# Structure and Cl^-^ Conductance Properties of the Open State of Human CFTR

**DOI:** 10.1101/2024.12.08.627282

**Authors:** Zhi-Wei Zeng, Christopher E. Ing, Régis Pomès

## Abstract

The cystic fibrosis transmembrane conductance regulator (CFTR) is an anion channel that plays a vital role in water and ion secretion on epithelial surfaces. Loss of function in CFTR causes the life-threatening disease cystic fibrosis (CF). The functionally open state of CFTR has so far eluded detailed structural characterization. Although multiple near-atomic resolution structures of CFTR have been solved under conditions that promote channel opening, they all lack a continuous ion conduction pathway. In recent molecular dynamics (MD) simulations, structural fluctuations of human CFTR in a hydrated lipid bilayer led to the observation of transient Cl^-^ conducting states, but the stability and conduction properties of these putative open states were not established. Here, we conduct massively repeated simulations initiated from these Cl^-^ permeable conformations. Reproducible structural relaxation of the pore leads to a stable open conformation featuring five symmetrically arranged pore-lining helices. Unlike previously reported structures, this novel penta-helical arrangement reproduces experimentally determined properties of the open pore, including a Cl^-^ conductance close to that measured at physiological voltages. Together, our results support the validity of this newly identified pore conformation as a model of the fully open channel. Detailed analysis highlights the role of cationic pore-lining residues in the Cl^-^ permeation mechanism and suggests that the kinks observed in several transmembrane helices play a role in channel gating.

## Introduction

CFTR is an anion channel that facilitates water and ion secretion from the cells of epithelial tissues (Berger et al., 1991; Riordan et al., 1989; Saint-Criq and Gray, 2017). It is vital for the healthy function of exocrine organs in respiratory (Smith et al., 1996), digestive (Ishiguro et al., 2009), and reproductive systems (Liu et al., 2012). Loss of function in CFTR causes cystic fibrosis, a devastating genetic disease that shortens life expectancy primarily due to its deleterious impact on lung health (Elborn, 2016; Smith et al., 1996). Over the past three decades, CFTR has been extensively studied with the aim of understanding the molecular pathology of cystic fibrosis – from its folding and processing (Bose et al., 2020; Lukacs and Verkman, 2012) to gating and ion conduction mechanisms (Hwang et al., 2018; Linsdell, 2017).

Even though CFTR functions as an ion channel, it belongs to the superfamily of ATP-binding cassette transporters – specifically, the ABCC exporter subfamily with type IV transmembrane fold (Dassa and Bouige, 2001; Ikuma and Welsh, 2000; Thomas et al., 2020). Like other ABCC proteins, CFTR consists of two transmembrane domains (TMDs), which form a gated pore in the membrane, and two nucleotide-binding domains (NBDs) which bind and hydrolyze ATP during gating. In addition, CFTR possesses a disordered R-domain that becomes phosphorylated by protein kinase A (PKA) before the channel undergoes ATP-dependent gating (Cheng et al., 1991). During gating, ATP binding promotes dimerization of the NBDs (Ikuma and Welsh, 2000; Levring et al., 2023); as a result, two ATP-binding sites are formed at the interface between the NBDs: a catalytic site and a degenerate site which are differentiated by their ability to hydrolyze ATP. Once it has been stabilized by bound ATP, the dimerization of NBDs promotes conformational changes in the TMDs and opening of the gate. Subsequent hydrolysis of ATP at the catalytic site ultimately results in the separation of the NBDs and closure of the gate.

Over the past decade, invaluable structural insights into the functional states of the gating cycle have been gained. Near-atomic resolution cryo-EM structures have been determined for the unphosphorylated, ATP-free state, as well as for the phosphorylated, ATP-bound state (Fiedorczuk and Chen, 2022a; Liu et al., 2019, 2017; Zhang et al., 2018, 2017; Zhang and Chen, 2016). The folded domains of CFTR can be regarded as consisting of two half-channels, each of which consists of one NBD, four TM helices from a TMD, and two crossover TM helices from the other TMD (Figure 1A) – namely, half-channel 1 (TM helices 1, 2, 3, 6, 10, 11 and NBD1) and half-channel 2 (TM helices 7, 8, 9, 12, 4, 5 and NBD2). Chloride ions enter the channel cavity through a lateral portal between the two half-channels. A bottleneck region, sometimes referred to in the literature as the channel gate (Gao and Hwang, 2015) or selectivity filter (Gao and Hwang, 2015; Levring and Chen, 2024; Linsdell, 2016), separates the lumen of the channel (inner vestibule) from the extracellular space (Figure 1B). The structures provide insights into the mechanism of channel activation via phosphorylation. Comparison of cryo-EM maps for unphosphorylated and phosphorylated CFTR revealed that, upon phosphorylation, the R-domain is disengaged from the two half-channels, allowing ATP-induced NBD dimerization to occur (Zhang et al., 2017; Zhang and Chen, 2016).

**Figure 1.**
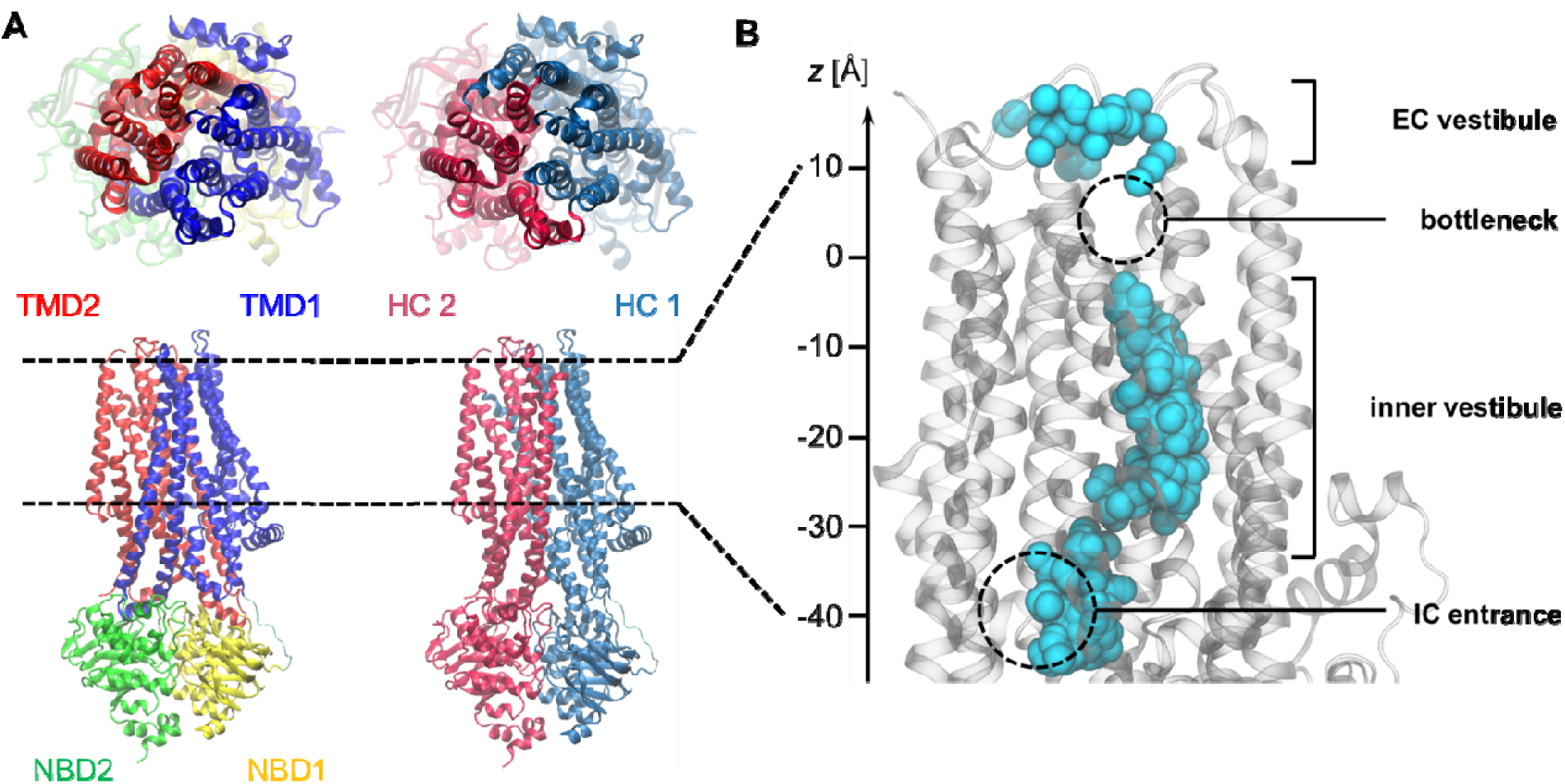
Overview of CFTR channel structure. **(A)** Cartoon representations of the 3D structure of CFTR (PDB: 6MSM). Different parts of the channel are coloured according to the domain (left) or the half channel (HC; right) to which they belong. The dashed lines indicate the membrane region. **(B)** Side view of the channel lumen. Cl^-^ ions (cyan spheres) from multiple simulation snapshots are overlaid to TM helices (gray cartoon), outlining the space accessible to ions inside the pore. Intracellular Cl^-^ ions cross a lateral intracellular (IC) entrance (*z*∼-40 Å) into the inner vestibule which leads to the pore bottleneck (*z*∼0 Å) separating the inner vestibule from its outer or extracellular (EC) counterpart. Note the absence of Cl^-^ from the bottleneck region in this “near-open” state of the channel.

Although more than ten near-atomic resolution (< 4 Å) cryo-EM structures of NBD-dimerized CFTR have been determined to this date (Fiedorczuk et al., 2024; Fiedorczuk and Chen, 2022a, 2022b; Liu et al., 2019; Zhang et al., 2018, 2017), none of them appear to provide an open pathway for Cl^-^ permeation – a defining feature of the open state. In these structures, the channel appears closed at the bottleneck despite its activation by phosphorylation, ATP and, in some cases, activity enhancing drugs (Fiedorczuk and Chen, 2022a; Liu et al., 2019; Zhang et al., 2018, 2017). Whether these phosphorylated, ATP-bound structures represent the functional open state has been elusive and somewhat contentious. On the one hand, some of these structures were proposed to represent the *bona fide* open state of the channel as they could narrowly accommodate fully dehydrated Cl^-^ ions in the gate region (Levring and Chen, 2024). However, this hypothesis disregards the high dehydration penalty of Cl^-^ and the fact that CFTR is also permeable to anions larger than Cl^-^, both of which considerations suggest that the pore bottleneck is significantly wider in the open state than in existing experimentally derived structures (Linsdell et al., 1997; Linsdell and Hanrahan, 1998). Accordingly, these ATP-bound structures have also been called “near-open” (Linsdell, 2018) or “quasi-open” (Simon and Csanády, 2021), implying that they closely resemble, but do not represent, the truly functional open state of CFTR. Uncertainties regarding the physiological relevance of structural models have been raised in studies of other ion channels (Dämgen and Biggin, 2020; Deng et al., 2020; Gonzalez-Gutierrez et al., 2017). In particular, there is evidence that the transmembrane solvent environment (lipids vs detergents) during structural determination can affect the structure of membrane proteins (Dalal et al., 2024; Deng et al., 2020). The use of detergents has been suggested to explain the observed closed-pore structures of CFTR despite the activating conditions (Cottrill et al., 2020; Zhang et al., 2017).

The above considerations raise the possibility that the near-open state may spontaneously relax to the open state in a lipid bilayer—a rationale that has been exploited successfully in MD simulation studies of other ion channels (Amaral et al., 2012; Cerdan et al., 2018; Chakrabarti et al., 2010; Shrivastava and Sansom, 2000) and motivated two recent computational studies of CFTR (Farkas et al., 2020; Zeng et al., 2023). In simulations of the near-open state of zebrafish CFTR, a structure highly similar to that of human CFTR, embedded in a lipid bilayer, the radius of the pore bottleneck was only wide enough to allow Cl^-^ passage transiently, with no spontaneous Cl^-^ permeation event observed on a sub-microsecond time scale (Farkas et al., 2020). Using an enhanced sampling technique to help Cl^-^ ions overcome the energy barrier in the bottleneck, two divergent Cl^-^ permeation pathways were identified (Farkas et al., 2020). In the other study, repeated μs-long simulations of the putative near-open state of human CFTR (Zeng et al., 2023) led to the observation of rare and transient Cl^-^ conducting bursts: in two out of ten simulations, a total of 17 Cl^-^ ions traversed the pore bottleneck in the presence of a strong TM voltage of -500 mV. Three distinct exit pathways were observed. Consistent with hydrophobic gating from a closed (dry or de-wetted) to a putative open (hydrated or wetted) state, these permeation bursts were accompanied by structural fluctuations of pore-lining TM helices leading to widening and wetting of the pore bottleneck.

These two simulation studies afford insight into the mechanism of Cl^-^ permeation that goes beyond the ion pathway inferred from static structures of CFTR (Farkas et al., 2020; Zeng et al., 2023). Consistent with the assumption that the latter are nearly open, the second study demonstrates that conformational fluctuations in the sub-microsecond time scale can lead to gate opening and ion permeation (Zeng et al., 2023). However, this result was obtained under an artificially high TM voltage, raising the question of whether the true open state of CFTR was attained. Indeed, the transient nature of the conduction bursts observed suggests that further structural relaxation may be required to reach a stable, fully open state that can support Cl^-^ currents at physiological voltages.

Here, we aim to elucidate the functional open state structure of human CFTR by building upon that previous study. To overcome sampling limitations, we performed hundreds of µs-long MD simulations extended from Cl^-^ permeable conformations of ATP-bound, NBD-dimerized human CFTR obtained previously. Over a total sampling time of 719 µs and across a broad range of voltage, further structural relaxation of the channel involving the reproducible rearrangement of helices in the outer region of the TMD led to a stable conformation featuring five symmetrically arranged pore-lining helices. This novel arrangement satisfies experimentally determined properties of the open pore, including Cl^-^ conductance, and likely represents a valid model of fully open human CFTR under physiological conditions. The Cl^-^ permeation mechanism and implications for channel gating are analyzed and discussed.

## Results & Discussion

### A novel, penta-helical pore conformation

We performed 719 replicates of microsecond-long unrestrained MD simulations of ATP-bound, NBD-dimerized human CFTR, most of which were initiated from Cl^-^ permeable conformations obtained in our previous study (Zeng et al., 2023). The overall structure of the protein remained stable, with an average RMSD < 2 Å for Cα atoms in the TM region (Figure 2 – figure supplement 1). The NBDs remained dimerized and the fluctuation of TM helices in the plane of the inner leaflet was limited (Figure 2B). In contrast, the amplitude of structural fluctuations in the plane of the outer leaflet, which hosts the extracellular end of the pore, was noticeably larger (Figure 2A). Remarkably, while the planar distribution of most TM helices relative to the rest of the helical bundle was essentially unimodal, both TM1 and TM11 sampled bimodal distributions, with one state closer to the TM helical bundle (“in”) and the other ∼4 Å farther away (“out”) (Figure 2A and C). The two states were sampled in nearly equal proportions by both helices (in:out ratio of 54:46 for TM1; 60:40 for TM11). Transitions between the two states of TM1 and TM11 were also well sampled, occurring in 240 and 328 out of 719 trajectories, respectively. Importantly, the positions of TM1 and 11 were anti-correlated, in that as TM1 moved away from the centre of the pore, TM11 moved towards it, resulting in a symmetrical pentagonal pore lined by TM helices 1, 6, 8, 11, and 12 (Figure 2C and D). The anti-correlation of TM1 and 11 positions is confirmed by principal component analysis (PCA) of the spatial coordinates of TM helices, which reveals that four conformational states, labelled α to δ, were sampled in our simulations (Figure 3A). The first principal component (PC1), which accounts for 25% of structural fluctuations in the conformational ensemble, captures the collective movements of TM helices 1, 2 and 11 leading to the symmetrical pentagonal pore or α state (Figure 3B and C, Figure 3 – figure supplement 2). In contrast, PC2, which accounts for 8% of structural fluctuations, captures comparatively more subtle movements of TM helices, as both β and γ states are characterized by a “squeezed” arrangement of pore-lining helices (Figure 3B and D). This squeezed arrangement also dominates in the δ state, which includes the near-open cryo-EM structure (PDB: 6MSM) and occupies a location intermediate between the other three conformational states (Figure 3A and B). To gauge the stability of the four conformational states, we analyzed the time evolution of their fractional population by considering all 719 MD trajectories in parallel. While the populations of α, β, and γ either increased or remained stable, the population of state δ decreased over the 1-µs simulation time, confirming that the cryo-EM structure is a metastable intermediate state in a lipid bilayer environment (Figure 3 – figure supplement 1).

**Figure 2.**
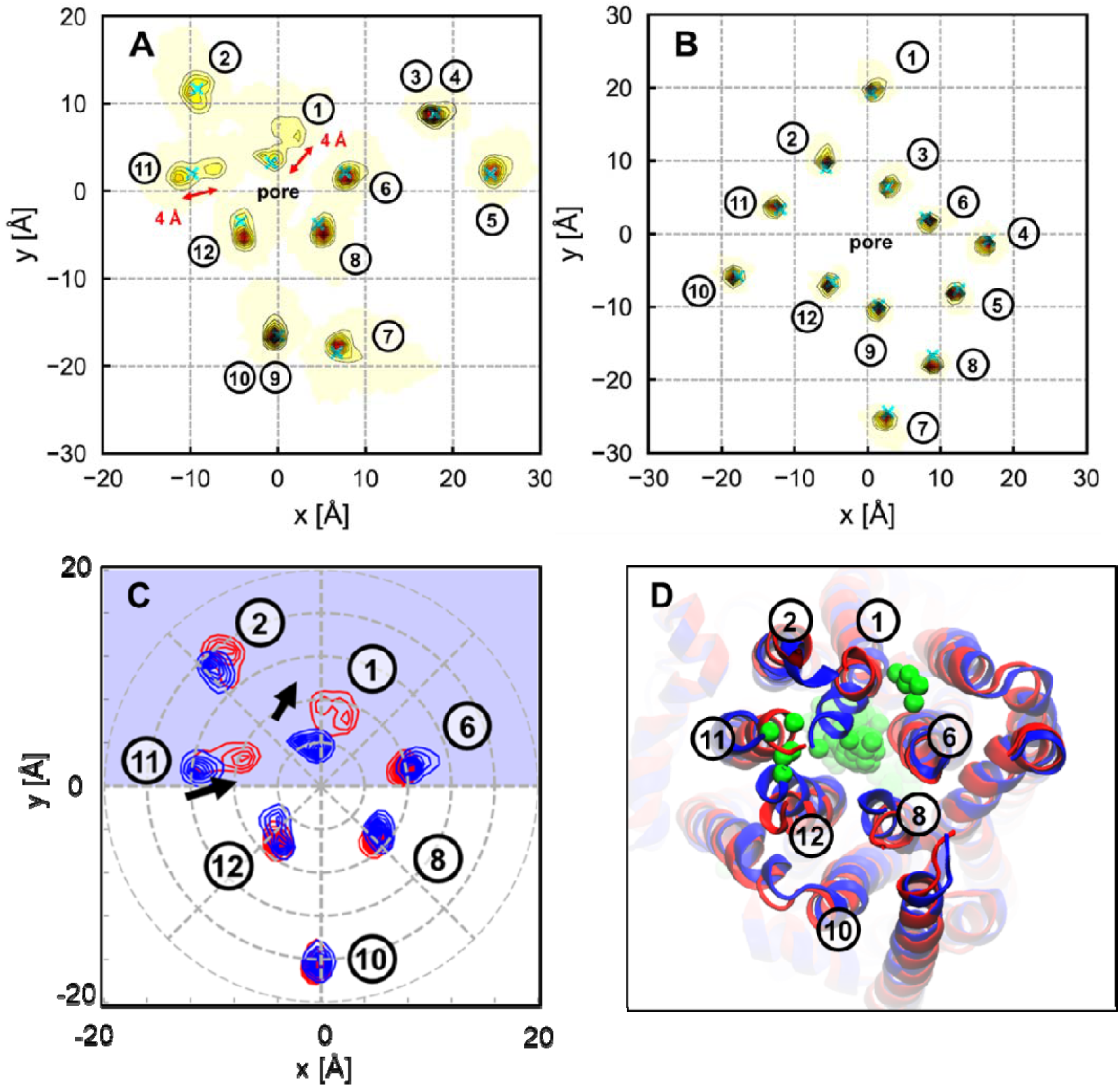
Conformational ensemble of TM helices. **(A)** Spatial distribution of TM helices in the plane (*x*,*y*) of the outer lipid leaflet. Conformational transitions result in well resolved bimodal distributions of TM helices 1 and 11. The extracellular ends of TM helices 3-4 and 9-10 overlap in this plane. **(B)** In contrast to the outer leaflet, the spatial distribution of TM helices in the plane of the inner membrane leaflet (*x*,*y*) is conserved throughout the simulations. **(C)** 2D histograms of outer leaflet positions of pore-lining helices (TM1, 6, 8, 11, and 12) and TM2 in the squished conformation of the pore (blue) versus expanded pentagonal conformation (red). Helices in the blue-shaded region belong to half-channel 1, whereas those in the unshaded region belong to half-channel 2. **(D)** Top (extracellular) view of representative structures of the squished (blue) and pentagonal (red) conformations superimposed. Cl^-^ ions (green spheres) are overlaid from multiple simulation snapshots to indicate the location of the channel pore.

**Figure 3.**
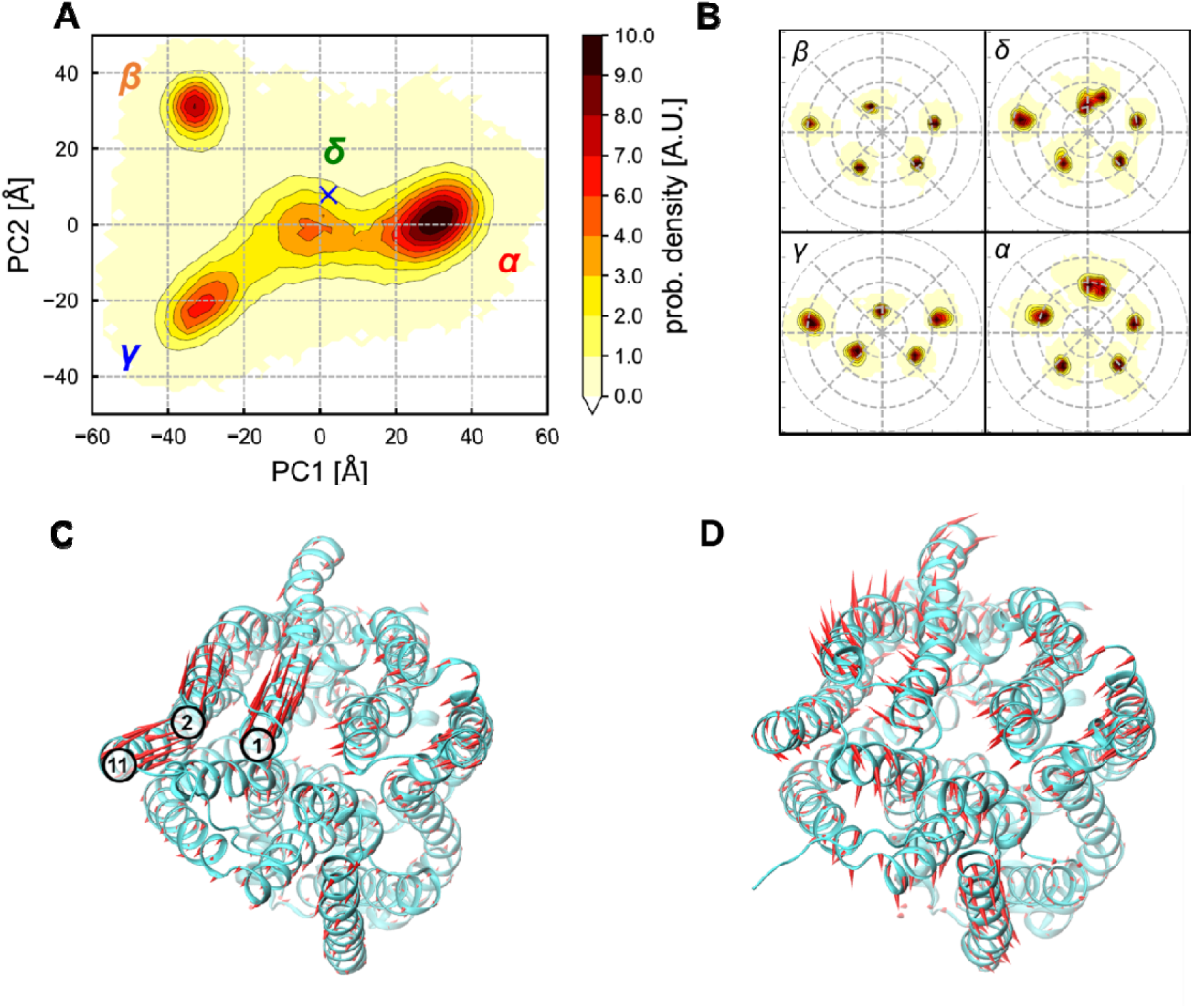
Principal component analysis of the CFTR conformational ensemble in the TM region. **(A)** Distribution of TM conformations projected onto the plane of the first two principal components reveals 4 conformational states labelled α-δ. The cross indicates the location of the initial cryo-EM structure (PDB: 6MSM). Contour levels are evenly spaced values of probability density. **(B)** Spatial distribution of pore-lining helices (TM1, 6, 8, 11, 12) in the plane of the outer leaflet (*x*,*y*) for conformational states α-δ. Contour levels are evenly spaced values of probability density. Concentric circles are centered on the pore axis and spaced by 4 Å. **(C**-**D)** Porcupine representation (red) of eigenvectors **(C)** PC1 and **(D)** PC2 overlaid on the CFTR structure (cyan) to illustrate the collective movements in each dimension. The largest collective motion, PC1, captures the concerted movement of TM1, 2, and 11 from near-open states β and γ to the pentagonal pore conformation, α (see also Fig. 2C).

Among all states sampled, the structure of state α is the most compatible with the open state of human CFTR. As a result of the symmetric arrangement of the five pore-lining helices, the radius of the pore bottleneck was larger by 0.5∼1 Å compared to the other states (Figure 4, Figure 4 – figure supplement 1). Notably, α is the only state featuring TM11 as a pore-lining helix due to the movement of this helix towards the centre of the pore. Such movement brings TM11 closer to TM6 and away from TM12, consistent with cross-linking experiments suggesting the close proximity of bottleneck-lining residues of TM6 and TM11 in the open state (Fig. 4 – figure supplement 2) (Wang and Linsdell, 2012). Furthermore, the solvent-accessible surface areas (SASA) of bottleneck-lining residues T1115 and S1118 from TM11 are both greater compared to states β-δ, consistent with experimental findings that these residues are accessible to aqueous reagents in the open state (Fig. 4 – figure supplement 2) (Fatehi and Linsdell, 2009). Overall, state α exhibits experimentally demonstrated features of the open pore, indicating that it most likely represents the open state of CFTR compared to the existing cryo-EM structures as well as all the other TM conformational states sampled in our simulations.

**Figure 4.**
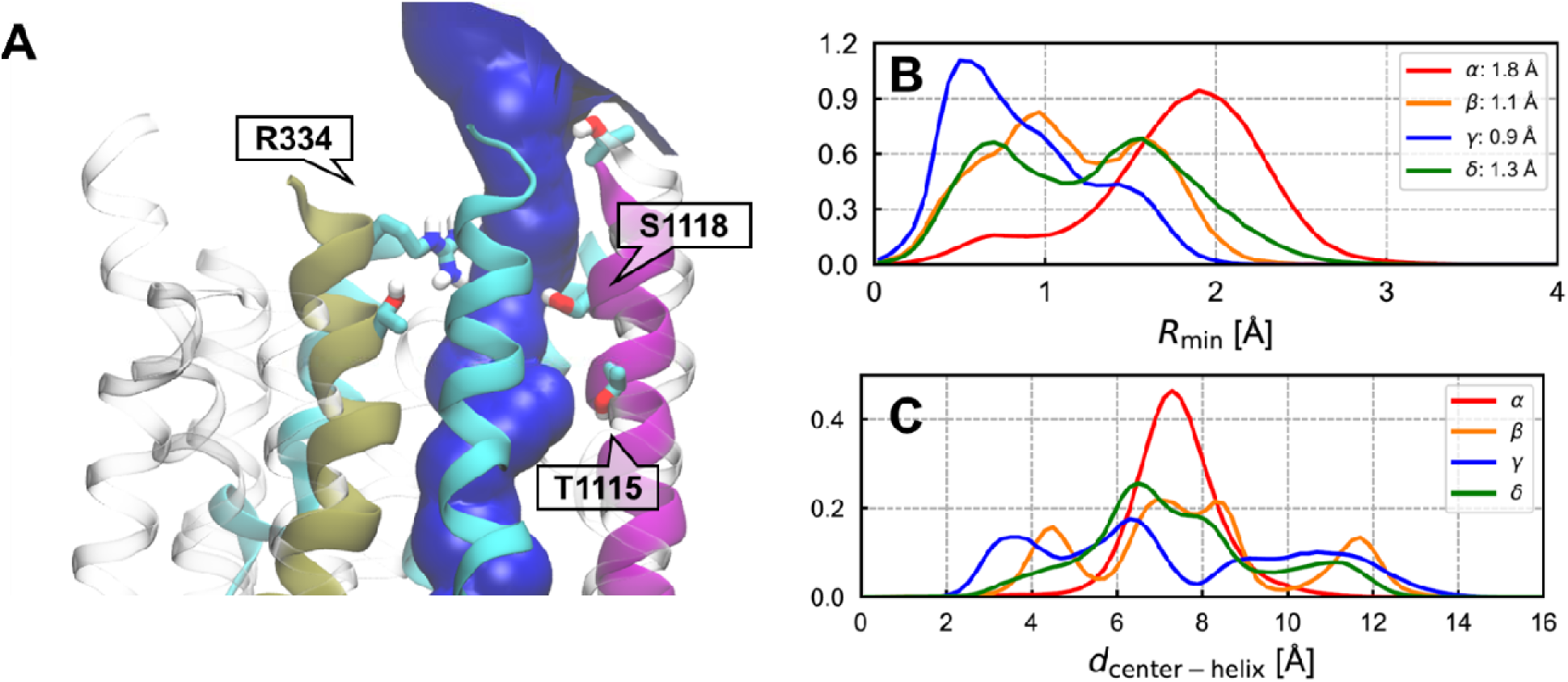
Structural features of the pore bottleneck. **(A)** Cartoon representation of the bottleneck region in putative open state α. Pore-lining helices are shown in cyan except for TM6 (tan) and TM11 (magenta). The blue surface encapsulating the interior of the open pore was computed with HOLE2 (Smart et al., 1996). **(B)** Distributions of the minimum radius (*R*_min_) in the pore bottleneck. Pore radii are calculated with HOLE2. Average values are indicated in the legend. **(C)** Distribution of the distances of pore-lining helices (TM1, 6, 8, 11, and 12) to the centre of the pore for conformational states α-δ. State α is unique for showing a unimodal distribution, consistent with the symmetrical arrangement of pore-lining helices.

### Cl^-^ permeation pathway

In our previous study of human CFTR, Cl^-^ permeation events occurred in 2 out of 10 microsecond-long MD simulations carried out at -500 mV TM voltage (Zeng et al., 2023). We identified three distinct Cl^-^ permeation pathways through the pore bottleneck: the “1-6” and “1-12” pathways, in which Cl^-^ ions exited through lateral openings between the extracellular ends of TM helices 1 & 6 and 1 & 12, respectively; and the “intermediate” pathway, in which Cl^-^ ions travelled along a straight path in between the other two pathways (Figure 5A and B). It was also found that the pathway taken by permeating Cl^-^ ions depended on minor differences in the structure and arrangement of pore-lining helices (Zeng et al., 2023). Remarkably, only one of these two simulations briefly sampled putative open state α. Moreover, in addition to α, Cl^-^ permeation events also occurred in states γ and δ, from which some Cl^-^ permeable snapshots served as starting configurations for the extended simulations in the present study (Figure 5 – figure supplement 1). Due to the small number of permeation events, the relative importance of the three permeation pathways as well as the corresponding pore structures could not be determined from the previous study. Our current simulation set contains more than 2000 Cl^-^ permeation events, amongst which all three permeation pathways were sampled. However, the pathway taken by Cl^-^ ions is strongly influenced by the conformation of the channel pore (Figure 5C and D). Whereas virtually all Cl^-^ permeation followed the 1-6 pathway in states β and γ, all three permeation pathways were sampled in both α and δ. Note that the enlarged pore bottleneck in state α allowed permeant Cl^-^ ions to occupy the region that previously separated the intermediate and 1-12 pathways, blurring the boundary between them (Figure 5A and B). As such, the intermediate and 1-12 pathways shall henceforth be collectively referred to as the “central” pathway.

**Figure 5.**
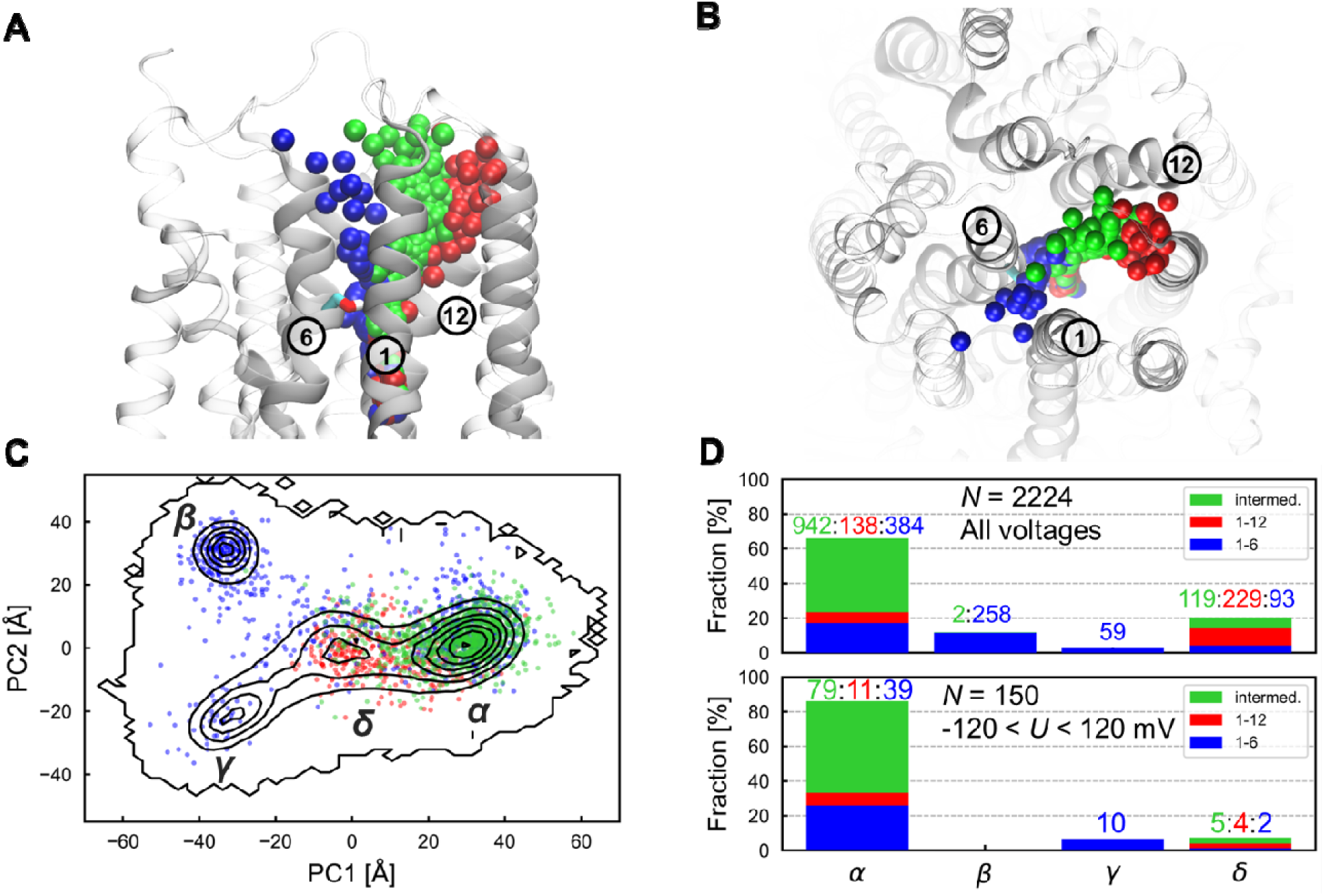
Cl^-^ permeation pathways through the pore bottleneck. **(A)** Side and **(B)** top views of the bottleneck region in the putative open state α with overlaid Cl^-^ ions (coloured spheres) showing three different Cl^-^ permeation routes. Blue: 1-6 pathway between TM1 and 6; red: 1-12 pathway between TM1 and 12; green: intermediate pathway. **(C)** Pore conformations in which Cl^-^ permeation occurs are indicated as scatter points on the (PC1,PC2) plane, coloured by permeation pathway. **(D)** Breakdown of Cl^-^ permeation pathways followed in each conformational state (*N*: number of permeation events) for (top) the full voltage range (-500 < *U* < 500 mV) and (bottom) the physiological voltage range (-120 < *U* < 120 mV). The number of times each permeation pathway was followed is indicated on top of the bar graphs.

Multiple Cl^-^ ion binding sites are present inside the pore. Many protein residues, primarily ones with polar and positively charged sidechains, engaged in direct contact with Cl^-^, displacing water molecules in its first solvation shell (Figure 6B and C). Consistent with our previous findings, there are two main Cl^-^ binding sites inside the channel cavity (Zeng et al., 2023): one near the intracellular entrance involving K190, R248, R303, R352, W356, and the other deeper inside the pore with R1097, K95, and R134 (Figure 6A). In addition, a third Cl^-^ binding site consisting of S341 and N1138 is found below the bottleneck.

**Figure 6.**
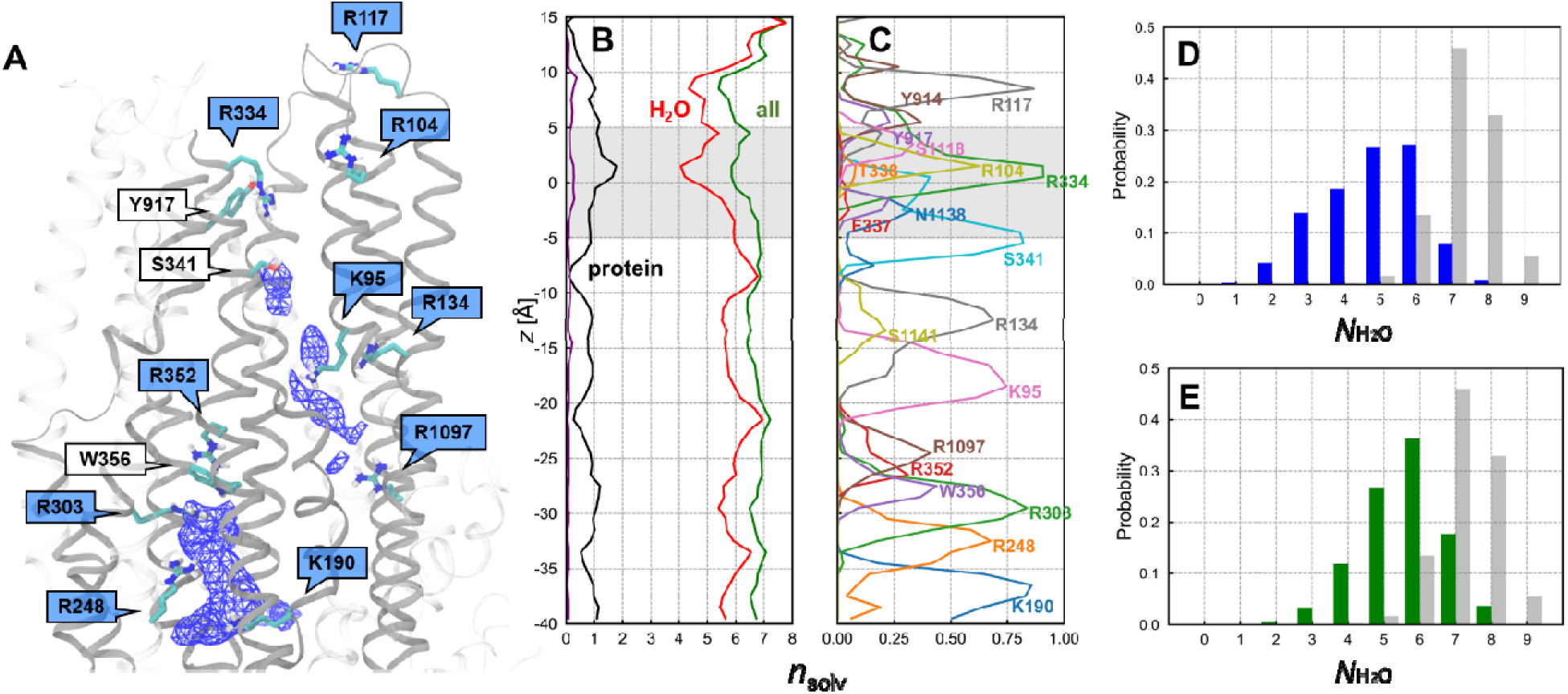
Solvation of Cl^-^ by protein and water inside the pore of CFTR. **(A)** Cl^-^ binding sites inside the inner vestibule. Regions of high Cl^-^ density (blue mesh) indicate the binding sites where Cl^-^ ions preferentially reside. Protein sidechains coming into direct contact with Cl^-^ ions in its first solvation shell are shown; blue labels highlight cationic residues among them. **(B-C)** Solvation environment of Cl^-^ ions that underwent translocation. The bottleneck region is indicated as the gray shaded region (-5 < *z* < 5 Å). **(B)** The average numbers of polar groups from protein sidechains (black) and of water molecules (red) in the first solvation shell of Cl^-^ ions (*n*_solv_) along the channel axis are shown together with their sum (green). The average number of non-polar protein residues in the first solvation shell of Cl^-^ ions is indicated in purple. **(C)** Axial distribution of protein residues making significant sidechain contributions to the solvation of Cl^-^ ions. F337 is also shown. **(D**-**E)** Distribution of the number of water molecules (*N*_H2O_) in the first solvation shell of Cl^-^ ions in the bottleneck region of **(D)** the 1-6 pathway (blue) and of **(E)** in the central pathway (green) compared to bulk water (gray).

Despite the abundance of hydrophobic residues in the bottleneck region (-5 < *z* < 5 Å), permeant Cl^-^ ions are partially solvated by polar and charged residues, including S341, T338, R334, Y914, Y917 and S1118 (Figure 6C, Figure 6 -- figure supplement 1). Direct binding to pore residues is responsible for the partial dehydration of Cl^-^ ions, which lose 1 to 3 water molecules out of an average of 7.3 ± 0.8 from their first solvation shell in bulk water (Figure 6D and E). Remarkably, in ∼15% of permeation events, the sidechains of non-polar residues lining the bottleneck, such as F337, also formed direct contacts with Cl^-^ ions and disrupted their solvation shell. Experimental studies have shown that mutations at residues R334, F337, T338, and Y914 affect the relative permeability among anions according to their hydration free energy – the greater the dehydration cost, the less permeable the ion, a mechanism known as lyotropic selectivity (Levring and Chen, 2024; Linsdell et al., 2000, 1998; Negoda et al., 2019). Taken together, these findings support the role of these residues in partial dehydration of Cl^-^ ions in the rate limiting step of permeation. Because these residues appear to line the 1-6 pathway in the cryo-EM structure, this pathway was recently suggested to be the only Cl^-^ permeation route (Levring and Chen, 2024). As noted above, it was proposed that the existing “near-open” cryo-EM structures in fact represent the functional open state of the channel, whereby fully dehydrated Cl^-^ ions pass through the 1-6 pathway without requiring further pore widening (Levring and Chen, 2024). Our findings challenge this proposed picture of Cl^-^ permeation in two respects. First, following TM movements, the bottleneck region of the pore expands, which allows Cl^-^ ions to permeate through two diverging pathways instead of a single pathway. Second, permeant Cl^-^ ions only ever undergo partial dehydration, regardless of which pathway (1-6 or central) is taken (Figure 6D and E, Figure 6 – figure supplement 1).

### Cl^-^ conductance

From numerous Cl^-^ permeation events at different TM voltages, we next evaluate the Cl^-^ conductance of the channel. To quantify the Cl^-^ conductance of states α–γ, we partitioned the simulation trajectories by state and computed the Cl^-^ current in terms of the number of translocation events over time spent in each state at different voltages. Across all negative TM voltages, the putative open state α is most permeable to Cl^-^ ions, exhibiting a Cl^-^ conductance of 3±1 pS, a value approaching the range of 6∼10 pS measured experimentally at physiological voltages (Sheppard and Welsh, 1999) (Figure 7). In contrast, states β-δ showed almost no Cl^-^ conductance at low negative TM voltages (-100 < *U* < 0 mV) and only produced Cl^-^ currents lower than state α at high negative voltages, suggesting that they are unlikely to be functional open states. At positive TM voltages, however, all four states virtually lost the ability to conduct Cl^-^ ions (e.g. 0.5±1 pS for state α at 0 < *U* < +100 mV). This result is at odds with the linear IV relationship across all voltages, from negative to positive (Sheppard and Welsh, 1999). Although some studies have reported minor inward rectification at positive voltages for the wild-type channel (Cai et al., 2003), the rectification of IV-curves computed from our MD simulations is much stronger.

**Figure 7.**
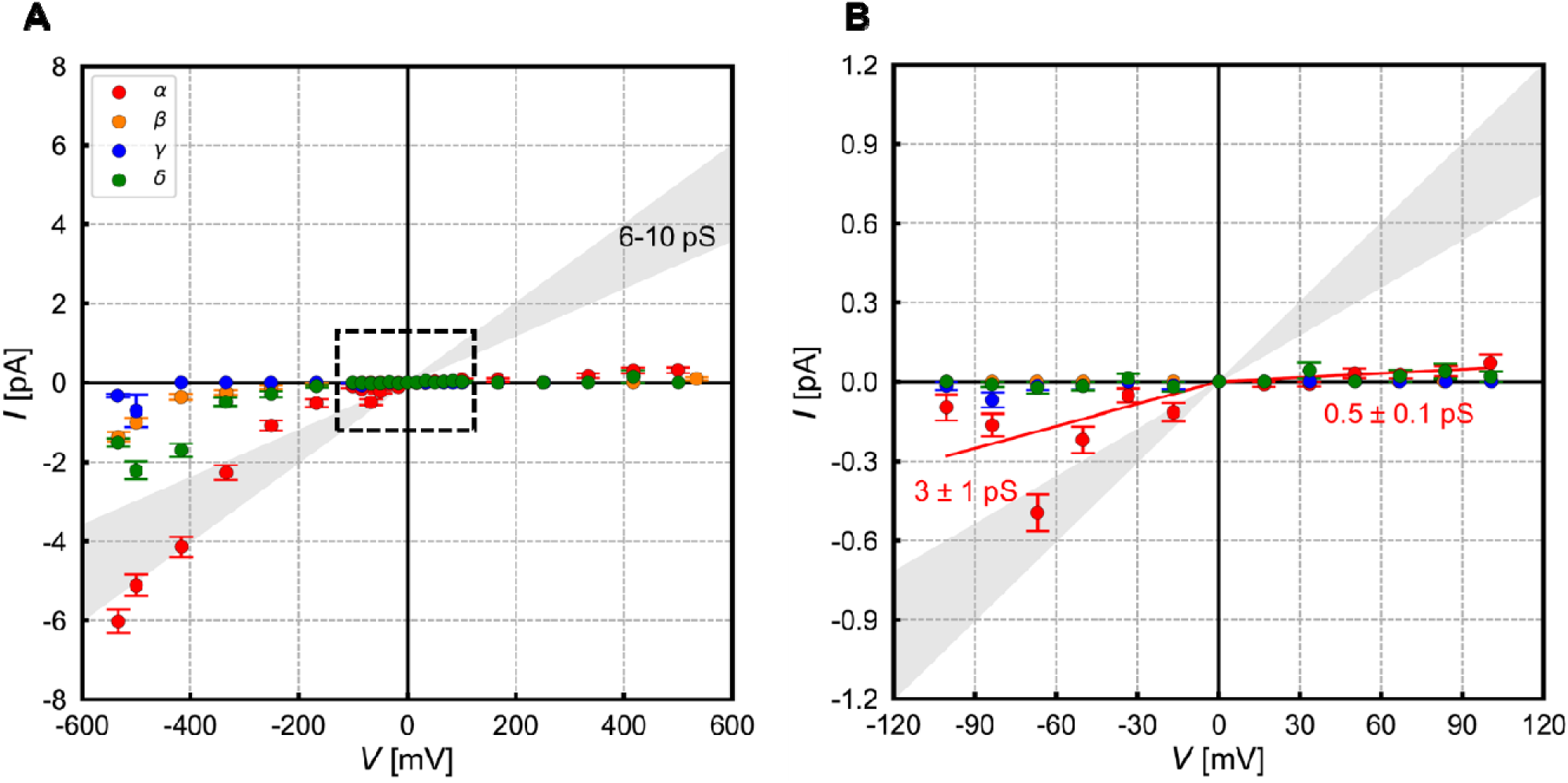
Current-voltage (I-V) relationship for each of the four conformational states. **(A)** Entire voltage range sampled and **(B)** physiological and experimentally accessible range. Zero-intercept linear regression fits of data points in the positive (0 < *U* < +120 mV) and in the negative (-120 < *U* < 0 mV) TM voltages are shown for state α as red straight lines with conductance (slope) indicated. The gray shaded regions are bound by the experimental single channel conductance range of 6-10 pS (Sheppard and Welsh, 1999). Negative and positive currents indicate Cl^-^ efflux to and influx from the extracellular space, respectively.

To explain the observed low conductance and strong inward rectification compared to experiment, we examined factors influencing Cl^-^ permeation. Chloride ions traversing the bottleneck region made frequent direct contact with positively charged R334. Notably, the sidechain of R334 can adopt two distinct conformations differing by 6 Å in the axial position of the center of charge: one pointing up and away from the pore bottleneck (“up”) as seen in the PDB structure; the other dunking down and extending into the bottleneck region (“down”) to form an ion pair with permeant Cl^-^ ions (Figure 8A and B). When R334 is down, the average Cl^-^ ion density in the bottleneck as well as Cl^-^ conductance are increased (Figure 8C-E). As a control, the Cl^-^ conductance of the channel was unaffected by the conformational state of the sidechain of R134, another pore-lining arginine from the inner vestibule whose sidechain sampled two slightly different conformations (Figure 8 – figure supplement 1), indicating that the observed conductance enhancement is unique to R334.

**Figure 8.**
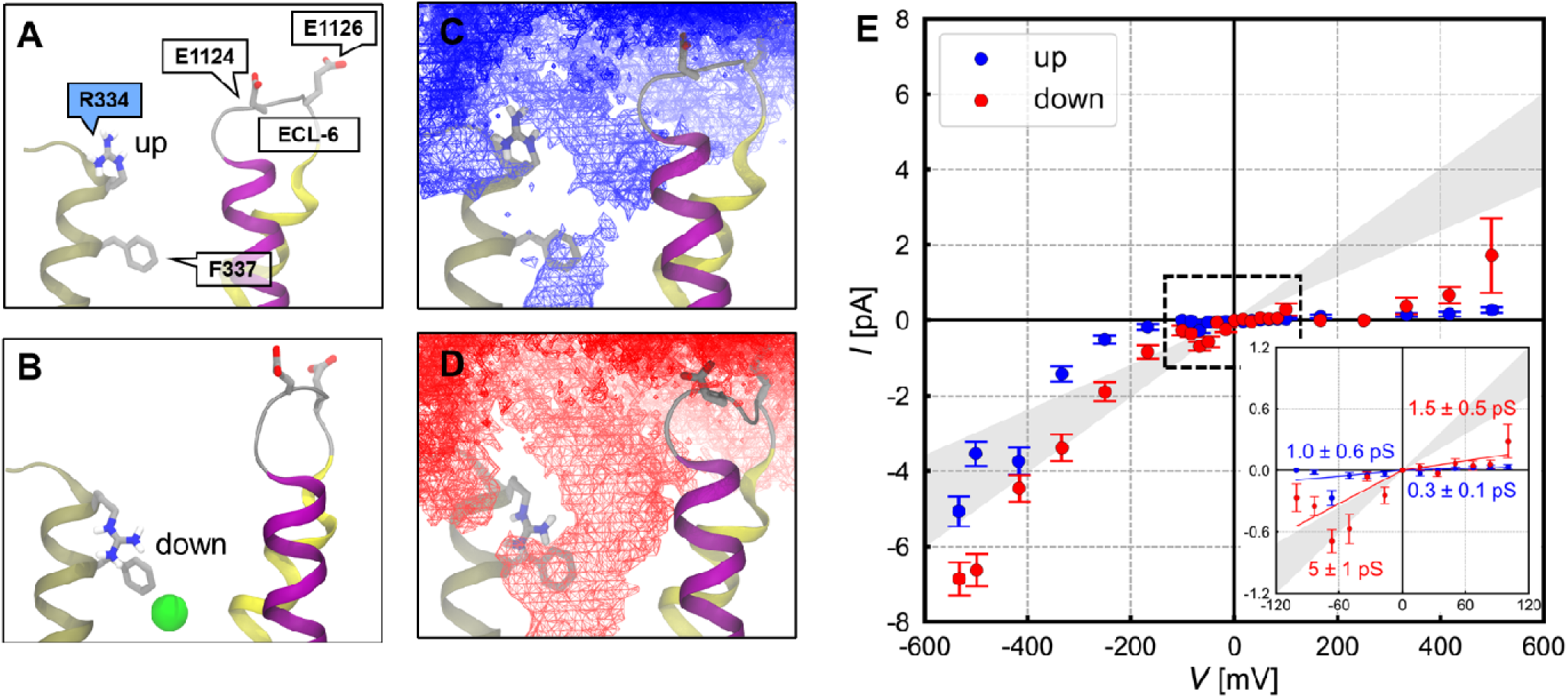
Effect of R334 conformation on Cl^-^ permeation. (A-B) Snapshots of the extracellular mouth of the pore with R334 in the **(A)** up and **(B)** down states. TM helices 6 (tan), 11 (magenta) and 12 (yellow) are shown together with loop ECL-6. In the down conformation, R334 can form cation-π and π-π interactions with F337 and an ion pair with Cl^-^ in the pore bottleneck. **(C-D)** Iso-surfaces reveal an increase in the average Cl^-^ ion density in the bottleneck region in state α from **(C)** the up state to **(D)** the down state of R334. **(E)** The current-voltage relationship shows that, in putative open state α, the down conformation of R334 (red) increases Cl^-^ conductance 5-fold in the physiological voltage range (inset) compared to the up conformation (blue).

Furthermore, the sidechain of R334, whether in the up or down conformation, can form a salt-bridge with either E1124 or E1126, two residues in the middle of flexible extracellular loop 6 (ECL-6) linking TM11 and 12. This interaction occurs when the loop is closer to the centre of the pore, resulting in the dunking of either glutamate sidechains (Figure 9A-B), blockage of the central ion permeation pathway (Figure 9E-F), and neutralization of the positive charge of R334 (Figure 9C-D). Glutamate dunking results in a 5-fold decrease in conductance in the range -100 < *U* < 100 mV in the penta-helical state (Figure 9G).

**Figure 9.**
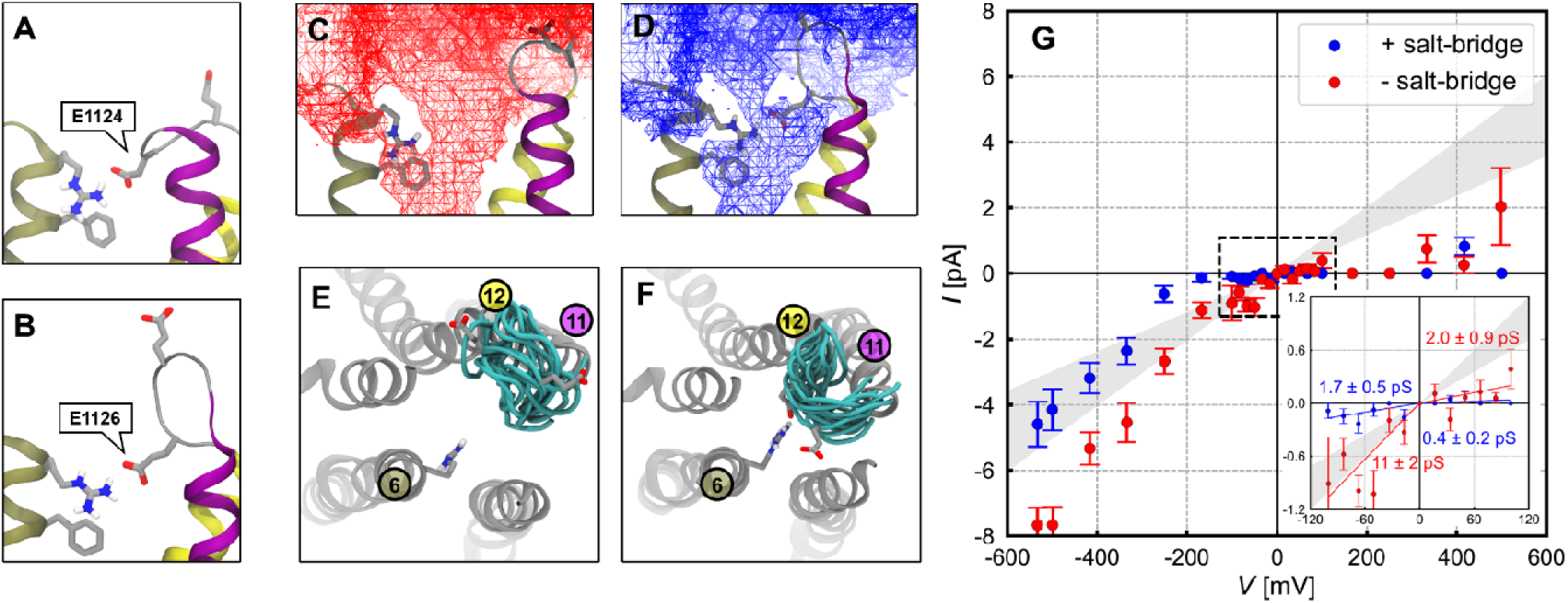
Effect of salt-bridge interactions between R334 and ECL-6 loop residues on Cl^-^ permeation. **(A-B)** Snapshots of the extracellular mouth of the pore showing the dunked sidechain of R334 forming salt bridges with **(A)** E1124 and **(B)** E1126. **(C-D)** Iso-surfaces reveal a greater Cl^-^ ion density in the extracellular mouth of the pore in open state α when **(C)** the glutamate sidechains are disengaged from the central pore, compared to **(D)** when they form a salt bridge with R334. **(E-F)** Top view of representative conformations of ECL-6 (cyan) in the **(E)** loop-outward and **(F)** loop-inward states. The helices lining the pentagonal pore are shown in gray. **(G)** The current-voltage relationship shows that the salt-bridge interactions decrease Cl^-^ conductance in the R334-down, open state of CFTR, especially in the physiological voltage range (inset).

Our analysis shows that the configuration of charged residues at the extracellular mouth of the pore affect the conductance of the penta-helical pore. The most favourable conditions for Cl^-^ conduction are: 1) the dunked conformation of R334, which assist Cl^-^ ions to cross the narrowest region of the pore; and 2) the disengagement of two ECL-6 Glu residues from the conduction pathway. The apparent conductance of the penta-helical pore at each voltage is strongly correlated with the fraction of time when these two conditions are met (Figure 9 – figure supplement 1). The fact that these conditions were seldom satisfied at positive voltages explains the observed rectification. This bias is likely due at least in part to the fact that the positive electric field tends to push cationic R334 upwards and anionic ECL-6 Glu sidechains downwards towards the pore, resulting in a greater likelihood of charge pairing between these residues. In addition, it is possible that initial conformations of the ECL-6 loop do not reflect equilibrium and more generally, that the conformational equilibria of the extracellular loops are not adequately sampled in the simulations. In particular, interactions with other extracellular loops that could help keep ECL-6 away and disengaged from the central pore may not be modelled accurately. In that perspective, the omission of 11 residues in glycosylated loop ECL-4 linking TM7 and 8, which was not modeled in structural studies to date or in the present study but is close to ECL-6, could alter the conformational properties ECL-6.

Experimental evidence supports the role of R334 in enhancing Cl^-^ conductance in CFTR. For instance, loss-of-charge, CF-causing mutation R334W shows reduced Cl^-^ conductance compared to wild-type (cftr2.org, n.d.; Sheppard et al., 1993). Our results suggest that the ability of R334 sidechain to extend into the pore bottleneck is an important feature of the functional open state of CFTR. In support of this finding, it was shown that R334 becomes less accessible from the extracellular space in the open state than in the closed state (Zhang et al., 2005). The down conformation of the R334 sidechain could be stabilized through cation-π and π-π interactions with the aromatic ring of F337 (Figure 8B, Figure 8 – figure supplement 2). Consistent with this observation, both R334K and F337A single-point mutants displayed reduced conductance (Ge et al., 2004; Gong and Linsdell, 2004; Linsdell et al., 2000). It should be noted that classical forcefields such as the one that we are using, which approximate electronic distributions as atomic point charges, do not explicitly capture the electric quadrupole contributions of aromatic rings in π-interactions (Lemkul, 2020; Mackerell, 2004; MacKerell et al., 1998). As such, it is possible that these interactions stabilize the down conformation of R334 sidechain to a greater extent than captured in our simulations, which could result in greater Cl^-^ conductance and a better agreement with experimental single channel recordings. On the other hand, there is no evidence supporting the roles of E1124 or E1126 in CFTR function. Instead, recent study suggests that E1124A and E1126P mutations retain or slightly improve the function of the channel (Simon and Csanády, 2021), suggesting that the presence of negative charges at these two positions is not required for function and may even be inhibitory, consistent with our conclusion.

Taken together, the above results validate state α as the functional open state of human CFTR. Although limitations were identified in the accuracy of the conformational ensemble of loop residues at the extracellular mouth of the pore, the pentameric conformation of pore helices derived in the current study is stable and consistent with CFTR function. Our findings provide a realistic, physiological depiction of Cl^-^ permeation through the open state of CFTR, which features a wide central pathway and a side pathway between TM helices 1 and 6, with a preference for the central pathway (Figure 5D). We shall henceforth refer to state α as the “MD-open” state. In contrast, ion translocation only occurs at large, non-physiological voltages for states β-δ, confirming their status as “near-open” conformations. States β and γ shall be referred to as stable “MD-closed” states to distinguish them from the metastable near-open state δ.

### Insights into the gating mechanism

Having characterized the structure of the functional open state of the channel, we next seek to gain structural insights into CFTR gating. In addition to extensive simulations of the NBD-dimerized, ATP-bound CFTR, we performed 20 replicates of 1-microsecond-long simulations of the ATP-unbound, NBD-separated cryo-EM structure of closed state human CFTR (PDB: 5UAK) in a hydrated lipid bilayer. Consistent with past MD simulation studies (Corradi et al., 2018; Tordai et al., 2017), the NBDs either partially or fully dimerized in 15 out of 20 trajectories (Figure 10 – figure supplement 1).

Dimerization occurred very rapidly due to the fact that the structural model is missing the R-domain that keeps two half-channels apart in the closed, unphosphorylated state of the channel. This omission makes the initial NBD-separated conformation highly unstable. In the presence of the R-domain, one expects that the dimerization of NBDs would occur much more slowly as the R-domain gradually disengages from the two half-channels upon phosphorylation.

Despite the rapid dimerization of the NBDs, the arrangement of the extracellular TM helical segments remained unchanged over the 1-µs simulation time (Figure 10 – figure supplement 2), indicating that further conformational changes in the TM region need to occur for the gate to open following NBD dimerization. Accordingly, comparison of the MD-open state with the structural ensemble obtained after fast NBD dimerization of the closed state highlights persistent structural differences in the extracellular segments of TM helices, especially in half-channel 2. In particular, pore opening requires movement of the extracellular end of TM8 towards the center of the pore by ∼8 Å and TM12 away from it by ∼7 Å (Figure 10A-C, Figure 10 – video 1). To accommodate the outward movement of TM12, the extracellular ends of TM helices 10 and 11 must move apart from each other (Figure 10D). The same conformational changes of TM helices 8 and 12 from the ATP-unbound closed state to the near-open state were previously identified based on the comparison between closed and near-open cryo-EM structures (Zhang et al., 2017). Contrary to the previous hypothesis, however, our results suggest that reaching the open state from near-open state does not involve further conformational changes in these two helices (Figure 2D).

**Figure 10.**
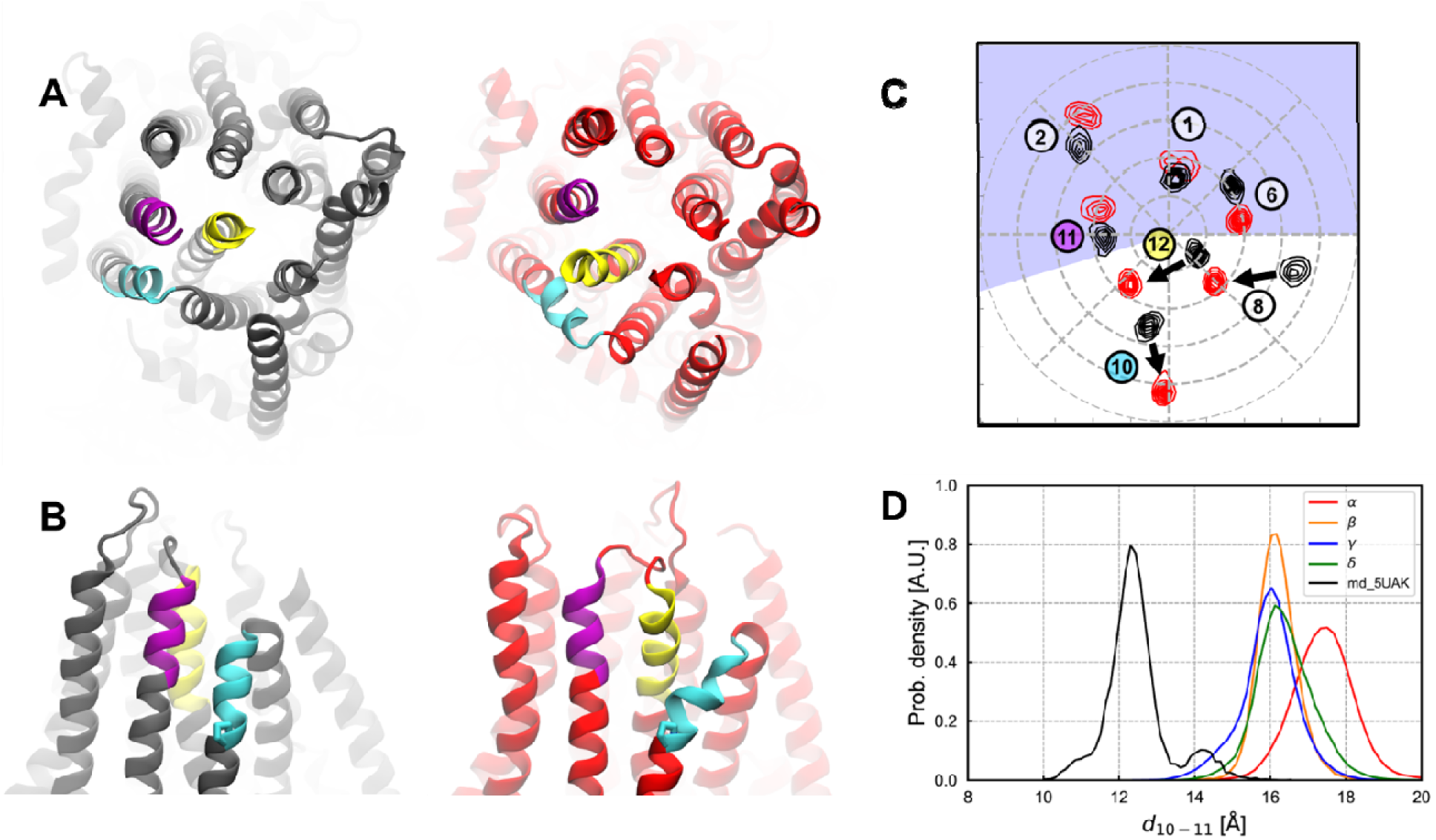
Structural differences between the ATP-unbound, closed state and the open state of human CFTR in the gate region. **(A)** Top and **(B)** side views of representative structures taken from MD simulations of ATP-free CFTR (left; black) and of the MD-open state of ATP-bound CFTR (right; red). To reach the open state, the outer segments of TM10 (cyan) and 11 (magenta) must separate to accommodate the intercalation of TM12 (yellow) between them. **(C)** 2D histograms of outer leaflet positions of pore-lining TM helices (1, 6, 8, 11, and 12) together with TM 2 and 10 from MD simulations of (black) ATP-free CFTR versus (red) MD-open state of CFTR. The largest movements of TM helices needed to reach the open state are indicated by arrows. Helices in the blue-shaded region belong to half-channel 1, whereas those in the unshaded region belong to half-channel 2. **(D)** Distribution of distances separating the centres of mass of the outer segments of TM10 and 11 in the ATP-free closed state (black) and in the four ATP-bound conformational states α-δ.

Whereas reaching the open state from the ATP-unbound, NBD-separated closed state primarily involves the closure of two half-channels at the NBDs and rearrangements of TM helices 8 and 12 within half-channel 2, reaching the MD-open state from near-open states primarily involves the movements of TM helices within half-channel 1, with the analysis above showing that collective motions of TM helices 1, 2, and 11 were the dominant changes involved in channel opening (Figures 2 and 3). What is the structural mechanism underlying such a movement? A notable aspect of the CFTR structure is the abundance of kinks in the TM helices, of which all except TM7 contain at least one kink. Helical kinks often result from structural distortions due to proline or glycine residues, which impart structural flexibility within the helix and have been suggested to facilitate conformational changes in other transmembrane proteins (Chamberlain et al., 2003; S.P. Sansom and Weinstein, 2000). In human CFTR, we identified five sites where helical kinks undergo changes between MD-closed and MD-open states: two on TM1, at G85 and P99; two on TM2, at G126 and P140; and one on TM11 at W1098 (Figure 11A). At P99, the average kink angle increases slightly from ∼20° in the MD-closed states to ∼25° in the MD-open state (Figure 11 – figure supplement 1A and B). In contrast, at G85, G126, and P140, reaching the MD-open state involves changes in the orientation of the kinks rather than in the magnitude of kink angles. Specifically, upon reaching the MD-open state from the MD-closed states: (1) the middle segment of TM1 (residues 85-96) swivels with respect to the intracellular segment (residues 78-85) by ∼20° (Figure 11F and G); (2) the extracellular segment of TM2 (residues 117-128) swivels with respect to the middle segment (residues 129-138) by ∼40° (Figure 11B and C); and (3) the middle segment of TM2 swivels with respect to the intracellular segment (residues 140-149) by ∼30° (Figure 11 – figure supplement 1C and D). Finally, at the W1098 hinge point, the kink angle is larger in the MD-open state by ∼8° compared to the MD-closed states, while the extracellular segment (residues 1098-1118) also swivels around the middle segment (residues 1077-1097) by ∼20° (Figure 11D). Among all observed patterns implicated for channel opening, the changes in kink at W1098 and in swivel at G85 and G126 are the most strongly correlated with the transition between MD-closed and MD-open states (Figure 11C, E and G). Altogether, changes in both the magnitude and the orientation of these helical kinks, even if not necessarily all occurring simultaneously, underlie the structural transition between MD-open and MD-closed states (Figure 11 – video 1).

**Figure 11.**
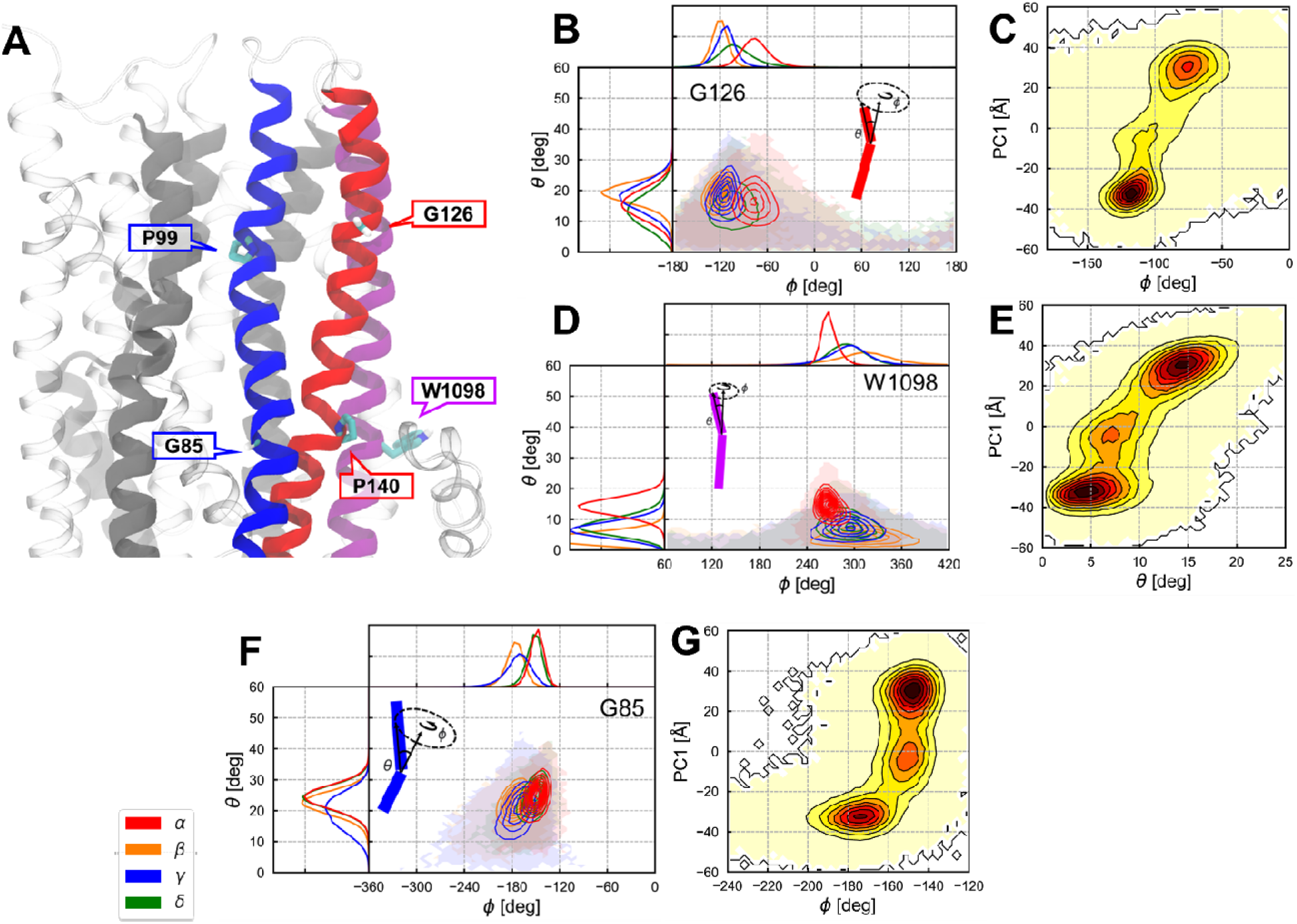
Analysis of changes in helical kinks underlying the transition between near-open and open states. **(A)** Cartoon representation of CFTR highlighting residues located at helical kinks on TM 1 (blue), 2 (red), and 11 (magenta) where structural changes occur upon channel opening. The sidechains of these residues (Hα atoms for glycine) are shown. The three helices completing the pore bottleneck are shown in dark gray. All other parts of the protein are translucent. **(B-G)** Analysis of helical kink magnitude and orientation. **(Left)** Distribution of kink angle θ (kink magnitude) and wobble or swivel angle (kink orientation) are shown for each of the four conformational states at **(B)** G126 on TM2; **(D)** W1098 on TM11; and **(F)** G85 on TM1. **(Right)** Contour plots showing the correlation between kink or swivel angles and principal component 1 (PC1), which captures conformational transitions between MD-closed and MD-open states (see Figure 3). Note that the uncertainty in kink orientation is high for small kink angles θ.

Studies of CFTR gating have understandably been focused on the coupling between conformational changes in the TMDs and ATP-binding-induced NBD dimerization. In this process, the gate opens in the TMDs in response to NBD dimerization, resulting in ion conduction bursts as observed in single channel recordings (Cai et al., 2011). Nevertheless, even during conduction bursts with dimerized NBDs, brief interruptions occur frequently (Cai et al., 2011; Levring et al., 2023). The transitions between conducting state and such “flicker closures” have been interpreted as ATP-independent gating events (Cai et al., 2003; Chen et al., 2017). Correspondingly, it is possible that conformational dynamic transitions such as those observed in our simulations of MD-open and MD-closed states correspond to such ATP-independent gating. In support of the functional relevance of helical kink flexibility at these sites, single-point mutations of G85, P99, and G126 are known to cause CF (cftr2.org, n.d.; Decaestecker et al., 2004; Sheppard et al., 1996; Wagner et al., 1994). Although functional and clinical data associated with P140 are not currently available, this residue is noteworthy for being in a π-bulge, a type of helical defect linked to functional roles in other proteins (Cooley et al., 2010). Finally, our analysis shows that changes in the magnitude of the kink angle near W1098 are strongly correlated to transitions between MD-closed and MD-open states (Figure 11E). This finding may seem surprising considering that unlike proline and glycine, tryptophan is not usually associated with helix kinks. However, consistent with the functional importance of this locus, W1098R and W1098C are CF-causing mutations (cftr2.org, n.d.).

Our results show that changes in helical kinks can affect the relative arrangement of pore-forming helices and are involved in near-open to open transitions in CFTR. Such changes are also likely to play a role in gating, as suggested by the fact that multiple residues located at helical kinks in TM helices are loci of CF mutations. The role of small changes in helical conformations in gate opening and closing has been noted in other ion channels. In the CorA magnesium channel, gating is coupled to a 10-degree change in the tilt of kinked pore-lining helices that modulates the diameter of the pore (Chakrabarti et al., 2010). In Kv and MthK channels, gating involves a ∼10-degree change in the internal kink of pore-lining helices (Fowler and Sansom, 2013; Gu and de Groot, 2020). In NavM channels, the open state is reached through a local α-to-π transition of a pore-lining helix (Choudhury et al., 2023; Choudhury and Delemotte, 2023). These examples, along with our findings on CFTR, show that the gating of ion channels can involve changes in helical structure as well as global conformational changes.

## Conclusions

In this study, we have derived and characterized a structural model of the open state of human CFTR through massively repeated MD simulations of a near-open structure of the ATP-bound, NBD-dimerized channel. Reproducible structural relaxation of the pore in a lipid bilayer leads to a stable open state structure featuring an approximately symmetric arrangement of five transmembrane helices lining an expanded pore bottleneck region. The functional relevance of this penta-helical open state is demonstrated by its ability to sustain Cl^-^ currents at physiological voltages with a conductance comparable to experimentally measured values. The detailed analysis of ion solvation and ion-channel interactions highlights the role of cationic residues in the ion conduction mechanism throughout the permeation pathway and reveals how the R334 sidechain facilitates the translocation of partly dehydrated Cl^-^ ions through the hydrophobic pore bottleneck.

While the present study does not address the allosteric mechanism linking NBD dimerization to channel opening, it provides insights into the gating mechanism in the pore domain (TMDs). The comparison of closed and open state structures suggests that ATP-dependent gating at long timescale primarily involves rearrangements of TM helices within half-channel 2, whereas the changes governing near-open to open transitions, which may be relevant to ATP-independent gating on a faster timescale, occur within half-channel 1. In this process, opening and closing of the hydrophobic pore bottleneck involves changes in the orientation and magnitude of flexible kinks in TM helices.

Understanding the structure and functional dynamics of CFTR is an important step towards unraveling the molecular pathology of CF. As a case in point, we note that several residues shown in this study to be implicated in pore opening and Cl^-^ permeation are loci of CF-causing mutations. Future studies of the atomistic open-state structure of human CFTR derived in this work should help gain a better understanding of gating and conduction as well as their defects in CF-causing mutants.

## Methods

### Molecular Systems

The preparation of the system containing ATP-bound, NBD-dimerized CFTR was described in detail in our previous work (Zeng et al., 2023). Briefly, the structural model of phosphorylated, ATP-bound human E1371Q CFTR (PDB: 6MSM) was used. The PDB structure is missing certain loop regions (residues 410-434, 890-899, 1174-1201, and 1452-1489) and the disordered R-domain (residues 638-844), which are not modelled in our study. All chains were acetylated at the N-termini and amidated at the C-termini. Two Mg-ATP moieties bound to the protein at the NBD interface in the PDB structure were kept, while all other putative lipids and an unknown helix at the TMD-NBD interface were removed. The Mg-ATP bound human CFTR was embedded in a hydrated POPC bilayer with 150 mM NaCl using CHARMM-GUI server (Jo et al., 2008).

For the ATP-free, NBD-separated CFTR, the simulation box was prepared in a similar way. The structural model of unphosphorylated, ATP-free human wild-type CFTR (PDB: 5UAK) was used (Liu et al., 2017). Residues that were missing in the PDB structure were not modeled, including a short N-terminal segment (1-4), the loops connecting each TMD-NBD pair (403-438, 1173-1206), the R-region (646-843), the extracellular loop between TM7 and TM8 (884-908), and the segment at the C-terminal end of NBD2 (1437-1489). A putative helix from the R-domain in the PDB structure with unknown amino acid composition was removed. Before embedding the CFTR protein into lipids, the PPM server of Orientations of Proteins in Membranes (OPM) database was used to determine the starting position of the lipid bilayer (Lomize et al., 2012). The model for CFTR with Mg-ATP bound was then embedded in a POPC bilayer and solvated in water with 150 mM NaCl using the CHARMM-GUI server without water or ions inside the channel pore (Jo et al., 2008). A total of 255 POPC molecules added around the protein per periodic box.

### Simulation setup and protocol

All MD simulations were conducted using GROMACS versions 2016 and later (Abraham et al., 2015). The CHARMM36 forcefield was used for protein, lipids, ions, ATP, together with the TIP3P water model (Best et al., 2012; Huang and MacKerell, 2013; Jorgensen et al., 1983; Klauda et al., 2010). Simulations were run at constant temperature and pressure (*T* = 300 K, *p* = 1 atm) at 2 fs integration timesteps. Constant temperature was maintained using the Nosé–Hoover thermostat (τ*_T_* = 0.5) (Hoover, 1985; Nosé, 1984); constant pressure was maintained using the Parrinello-Rahman barostat (τ_p_ = 2.0) (Nosé and Klein, 1983; Parrinello and Rahman, 1980). Semi-isotropic pressure coupling was used, with isothermal compressibility set to 4.5×10^-5^ bar^-1^ both in the *xy*-plane and along the *z*-axis. Nonbonded interactions were calculated using Verlet neighbor lists (Páll and Hess, 2013; Verlet, 1967). Lennard-Jones interactions were cut off at 1.2 nm and a force-based switching function with a range of 1.0 nm was used. The particle-mesh Ewald (PME) method was used to compute electrostatic interactions with a real-space cut-off of 1.2 nm (Darden et al., 1993; Essmann et al., 1995). The LINCS algorithm was used to constrain covalent bonds involving H atoms (Hess, 2008).

For initial simulations started from the PDB structures of CFTR, the system was subjected to energy minimization followed by a multi-step *NpT* equilibration protocol. The system was first subjected to steepest descent energy minimization until maximum force dropped below 1000 kJ/mol/nm. Random velocities were generated at the beginning of the *NpT* equilibration phase, which was conducted in three 10-ns stages, successively with protein heavy atoms, protein backbone atoms, and protein Cα atoms restrained (force constant *k* = 1000 kJ/mol/nm^2^ in *xyz* directions). *NpT* equilibration was followed by production run without restraints. All production run was 1-μs-long. For simulations with TM voltage, a constant electric field of strength between -32 mV/nm and +32 mV/nm was applied along the *z*-axis of the simulation box which is normal to the lipid bilayer. During the production runs, the *z*-dimension of the simulation box stabilized to and fluctuated around 16.5 nm. In all simulations, new random velocities were generated at the beginning of production runs. For extended simulations, snapshots from different timepoints of other MD trajectories were used as the initial configurations. No extra energy minimization or equilibration phase precedes extended production runs.

Among all simulations incorporated in the analysis, 649 of which started from both Cl^-^ permeable and impermeable configurations obtained in our previous study, and 50 of which were further extended from these simulation trajectories. For completeness, the dataset analyzed herein also includes 20 simulation trajectories obtained from our previous study, which started from the experimental “near-open” structure (PDB: 6MSM). Information about the initial configurations and how the extended trajectories are spawned are presented in Table 1 and Supplementary file 1.

### General analysis methodology of simulation data

Prior to analysis and visualization, CFTR structure from all snapshots of production run were aligned through minimization of RMSD of Cα atoms of TMDs. After alignment, the overall permeation pathway is parallel to the *z*-axis, which was used as the channel pore axis. The origin of the z-axis was approximately at the level of T338, and the POPC bilayer spanned the range from -30 Å to 10 Å on the z-axis. All visualizations and renderings of molecules were done with VMD (Humphrey et al., 1996). Scripts written in Tcl and python were used to perform analyses.

### Analysis of conformational ensemble of TM helices

The *xy*-positions of TM-helices were represented by the Cα of residues that lie approximately at the same *z*-level. At the inner leaflet level, Cα of residues 82, 142, 197, 241, 303, 356, 863, 938, 990, 1033, 1097, and 1152 were used to represent TM helices 1-12. At the outer leaflet level, Cα of residues 106, 121, 219, 324, 334, 882, 914, 1011, 1118, and 1131 were used. Due to that TM3, 4, 9, and 10 end at this level extracellularly, residues 219 and 1011 were used to represent the positions of helix pairs TM3-4 and TM9-10, respectively.

### Principal component analysis

Principal component analysis (PCA) was performed on the 3D Cartesian coordinates of TM helical backbone atoms (i.e. carbonyl carbon and oxygen, α-carbon, and amide nitrogen atoms) from all production run snapshots. K-means algorithm was used to cluster TM conformational states of CFTR. Both PCA and clustering were aided by Scikit-learn python library (Pedregosa et al., 2011).

### Pore dimension analysis

Pore dimensions were analyzed by HOLE2 program implemented through MDAnalysis (Gowers et al., 2016; Michaud-Agrawal et al., 2011; Smart et al., 1996). Residues 1-380 and 846-1173, including all hydrogens, were used to define the channel for HOLE2 analysis. The radius of the bottleneck in a given snapshot is represented as the minimum of the pore radius in the region -5 < *z* < 5 Å computed by HOLE2. Other parameters for running HOLE2 are included in the associated repository.

### Solvent accessibility analysis

Calculation of solvent accessible surface areas (SASAs) for residues T1115 and S1118 using VMD *measure* functionality (Humphrey et al., 1996). A probe radius of 1.4 Å was used, which corresponds to a coarse-grained spherical model for water. As T1115 is located at the protein-lipid interface, both protein and POPC lipids contribute to the solvent-excluded region during calculation.

### Analysis of Cl^-^ translocation and coordination

A Cl^-^ permeation/translocation event occurred if it: 1) crossed through *z* = 0 in the middle of the pore bottleneck; and 2) left the pore region subsequently without re-crossing *z* = 0. The time of translocation is when crossing *z* = 0 occurs. Details about the pore region definition are provided in the associated repository.

Time series for the 3D positions of the 17 Cl^-^ ions that underwent translocation from our previous study (Zeng et al., 2023) were grouped into three clusters based on similarity measured by dynamic time warping (DTW) distance (Sakoe and Chiba, 1978). The three clusters were visually verified to correctly correspond to the 1-6, intermediate, and 1-12 pathways. To assign translocation paths to all Cl^-^ permeation events in a high throughput manner, all trajectories of permeating Cl^-^ ions were collected and clustered into three categories based on their DTW distance to the cluster centroids. DTW analysis and clustering were aided by Tslearn python library (Tavenard et al., 2020).

Analysis of binding and solvation of Cl^-^ ions were aided by MDAnalysis. A molecule or residue is considered to form a direct contact with a Cl^-^ ion if it contains atom within 3 Å of the Cl^-^ ion, based on radial pair distribution functions computed in our previous study (Zeng et al., 2023). Low voltage simulations (i.e. -120 mV ∼ +120 mV) were used to analyze Cl^-^ binding and solvation.

### Analysis of current-voltage relationships

Transmembrane voltages *U* were computed by U= E_Z_ L_Z_, where *Ez* is the electric field strength in the *z*-direction, and *Lz* is the length of the periodic simulation box in the *z*-direction (Gumbart et al., 2012; Roux, 2008). At each given TM voltage, the simulation dataset was partitioned by the states of CFTR (i.e. TM helix conformation state α-δ, arginine sidechain conformational state, or ECL-6 conformational state). Chloride current at a certain voltage in a certain state is computed by *I* = *eN*/*t*, where *e* is the elementary charge, *N* is the number of translocation events, and *t* is the total time spent in the state. Simple linear regression without intercept was used to construct the current-voltage curve and determine the channel conductance (i.e. slope of the fit to current vs voltage plot).

### Helix kink analysis

The descriptors used to characterize helix kink, specifically kink and wobble/swivel angles, are inspired by the ProKink approach (Visiers et al., 2000). The kink angle between two helical segments were calculated as the angle between their principal axes. To compute the swivel angle, the protein structure from all snapshots were reoriented such that one of the helical segments is aligned with the *z*-axis. The swivel angle is represented as the azimuth angle of the other helical segment around the z-axis.

## Supporting information

trajectories_info

## Acknowledgements

We thank Paul Linsdell and Christine E. Bear for discussion of early results. This work was supported by Canadian Institutes of Health Research Grant MOP130461. MD simulations and analyses were enabled by supercomputing resources and support provided by SciNet (www.scinet.ca) and the Digital Research Alliance of Canada (www.alliancecan.ca). Z-W.Z. is supported by SickKids Restracomp.

## Data availability

Data generated from the study is provided on Zenodo at 10.5281/zenodo.14323352. Code created to analyze the data is provided on Github at https://github.com/wilzzw/cftr2.

## Author contributions

Z-W.Z. and R.P. conceptualized the study; Z.W.Z. and C.E.I. performed the MD simulations; Z.W.Z. analyzed the data; Z.W.Z. and R.P. wrote and edited the manuscript.

---

**Figure 2 – figure supplement 1.**
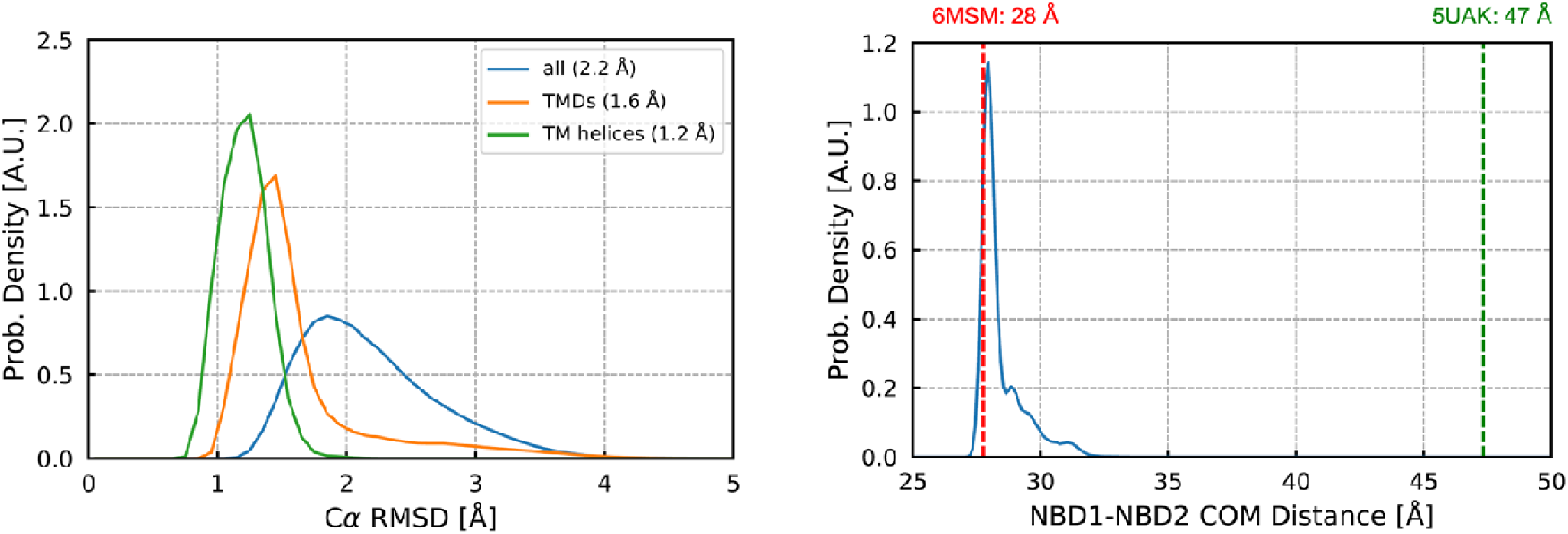
Conservation of overall structure of ATP-bound CFTR in MD simulations. Left: Distributions of RMSDs from the average of the MD structural ensemble. Average values are indicated in the legend. Right: Distribution of the distance between the centres of mass of the two NBDs (blue). The distances in experimental structures of ATP-bound (PDB: 6MSM; red dashed line) and ATP-free (PDB: 5UAK; green dashed line) states are shown for comparison.

**Figure 3 – figure supplement 1.**
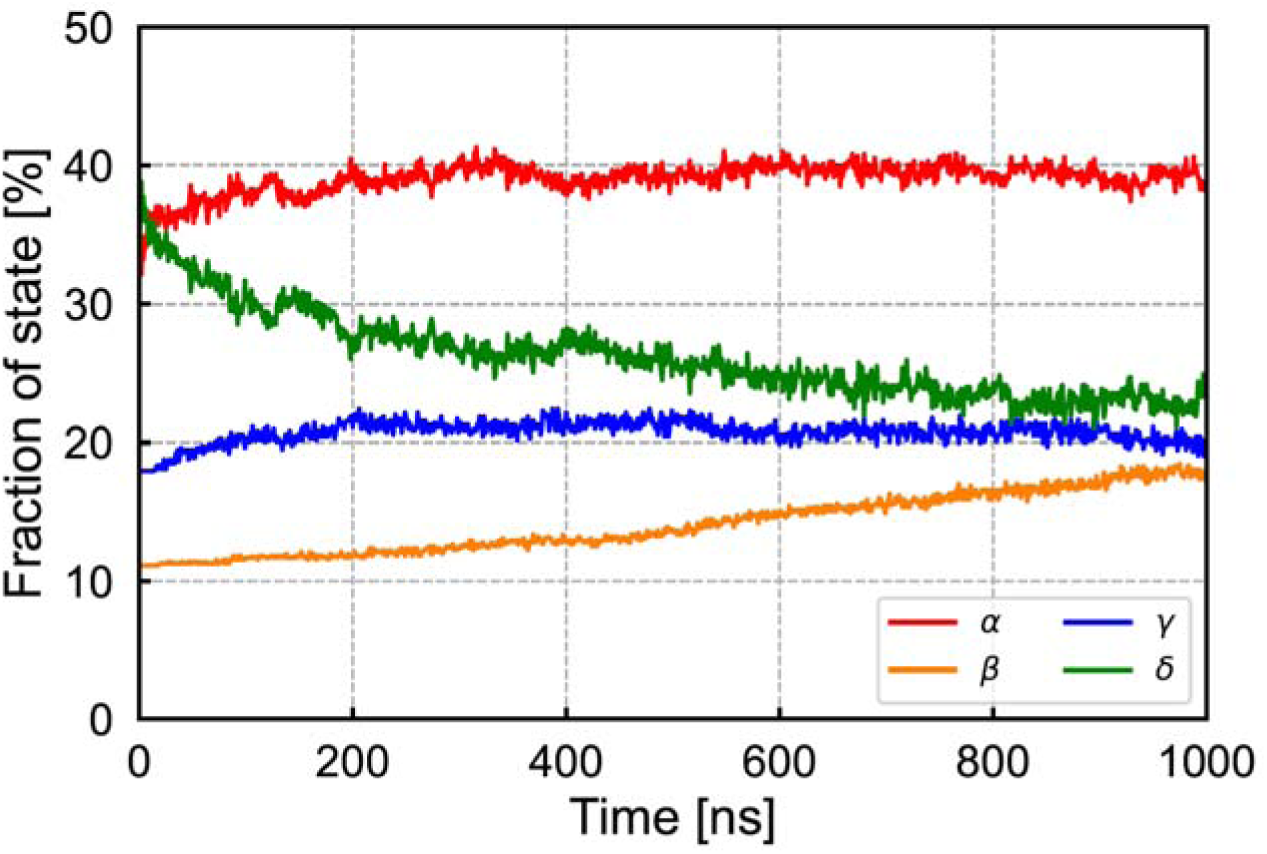
Time evolution of the fractional population of conformational states α-δ indicating their relative stability on the timescale of the MD simulations.

**Figure 3 – figure supplement 2.**
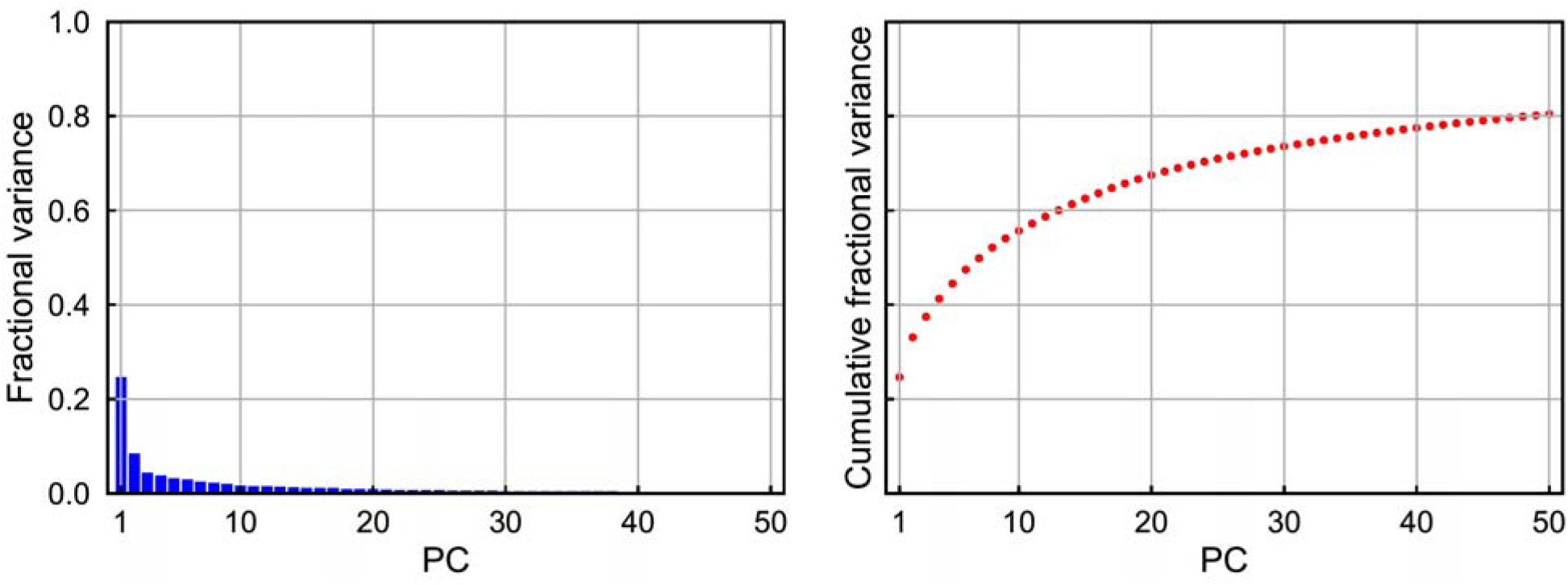
Supplementary information for PCA analysis. Left: fractional variance explained by each component. Right: cumulative fractional variance of principal components.

**Figure 4 – figure supplement 1.**
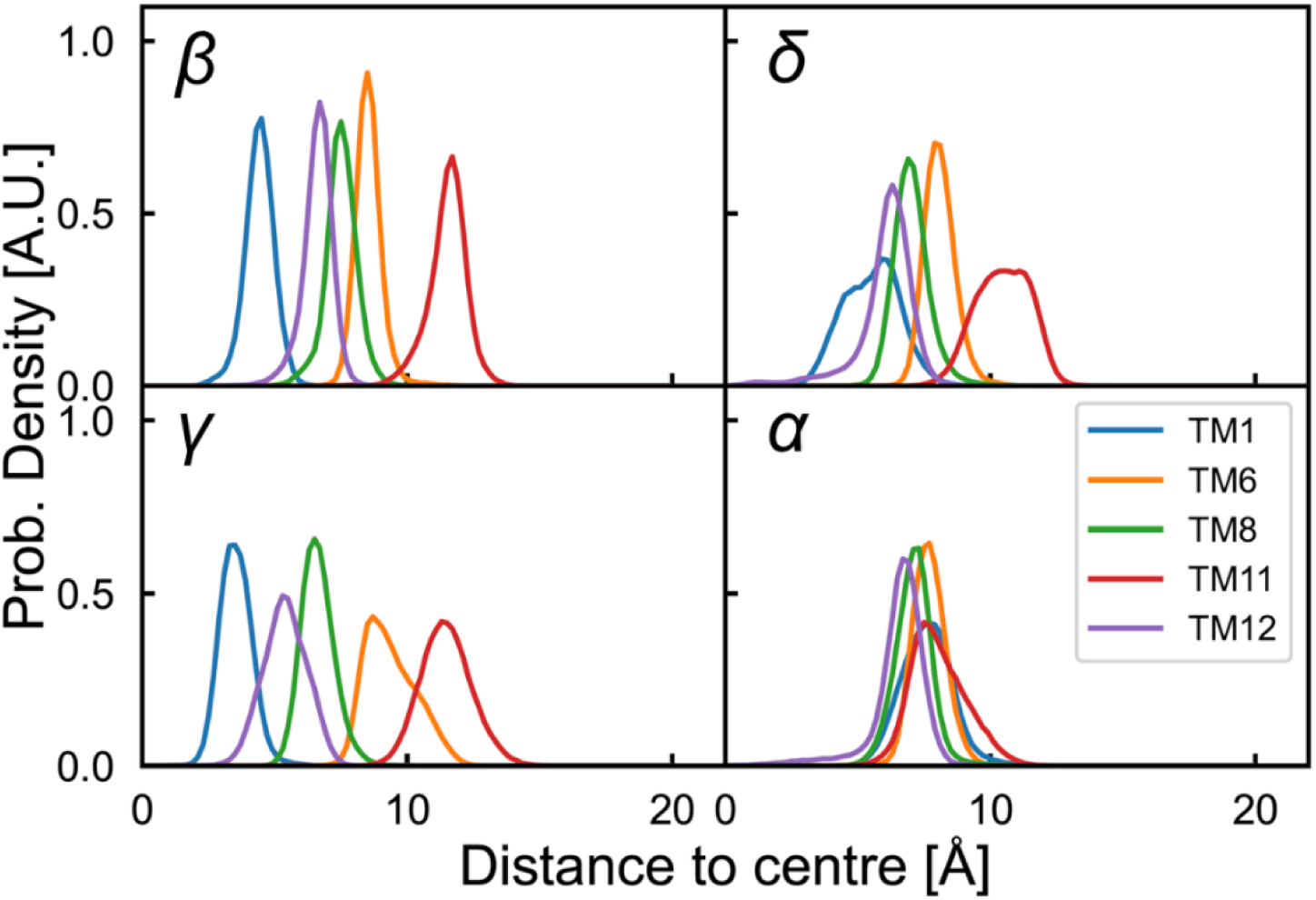
Symmetry of TM helix conformations in the pore bottleneck. Distributions of the distances of the five pore-lining helices to the centre of the pore in conformational states α-δ. In contrast to states β-δ, the strong overlap between narrow TM helix distributions in state α reflects their nearly symmetrical arrangement.

**Figure 4 – figure supplement 2.**
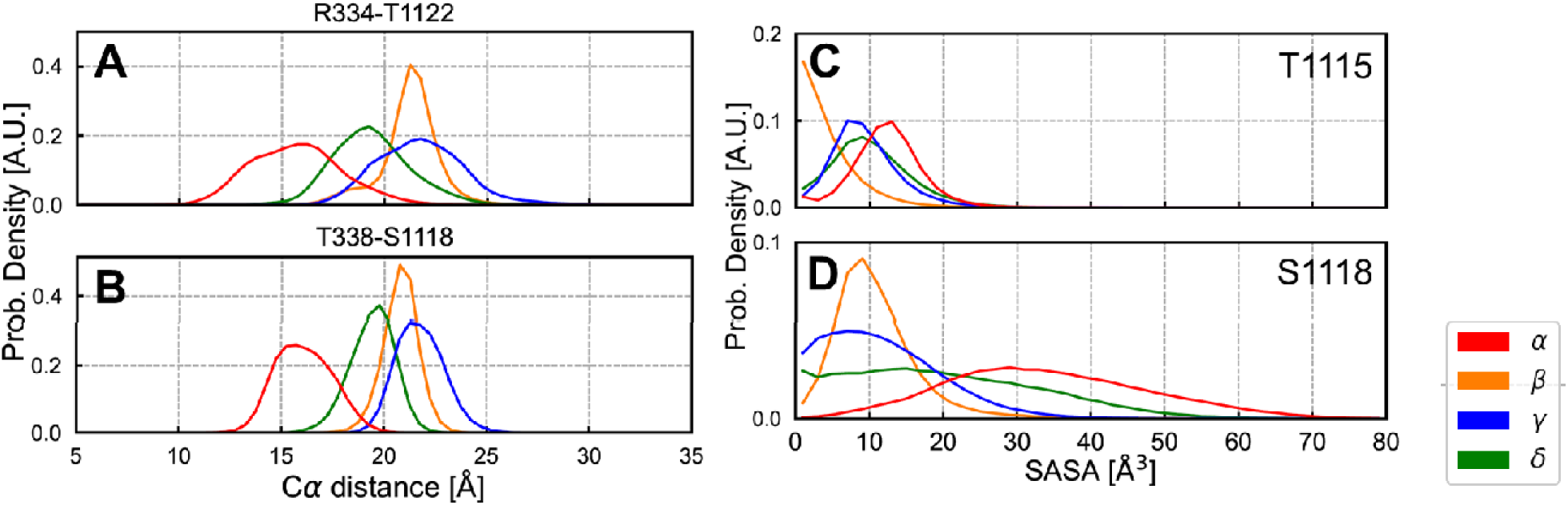
Detailed structural features of the pore bottleneck. **(A-B)** Distributions of distance between the Cα of residues R334 and T338 onTM6, and that of T1122 and S1118 on TM11 in the bottleneck region for conformational states α-δ. **(C-D)** Distributions of solvent accessible surface area (SASA) of bottleneck-lining residues T1115 and S1118 from TM11.

**Figure 5 – figure supplement 1.**
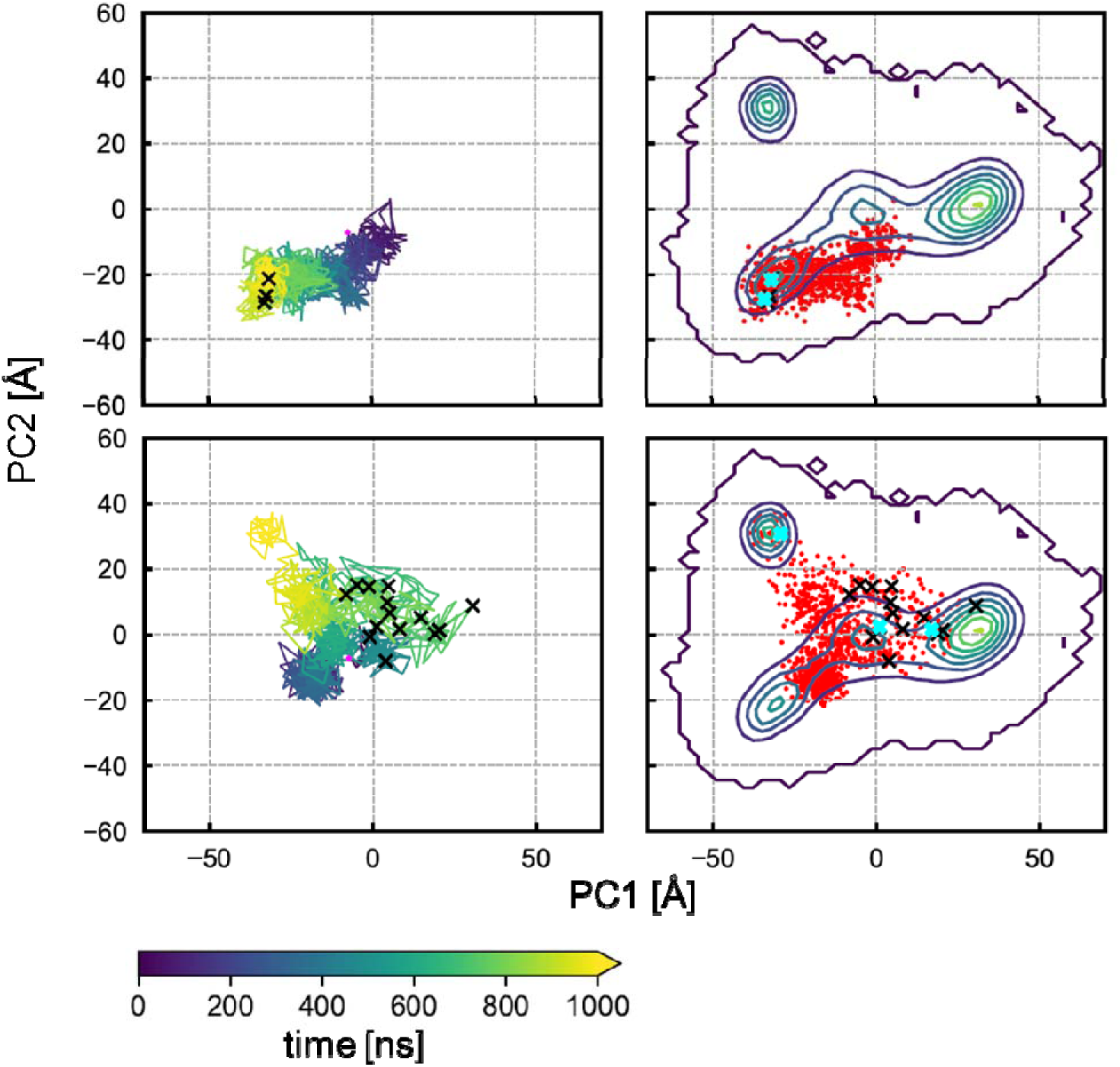
Projection of the two earlier trajectories in which Cl^-^ permeation events occurred in our previous study (Zeng et al., 2023) onto the (PC1,PC2) plane. **(Left)** traces of the two trajectories coloured by timestep. **(Right)** The conformations sampled in the two simulations indicated as red scatter points. Conformations in which Cl^-^ translocations occurred are marked in black “x”. The snapshots from these two trajectories used to initiate the extended trajectories reported in this work are marked in cyan “x”.

**Figure 6 – figure supplement 1.**
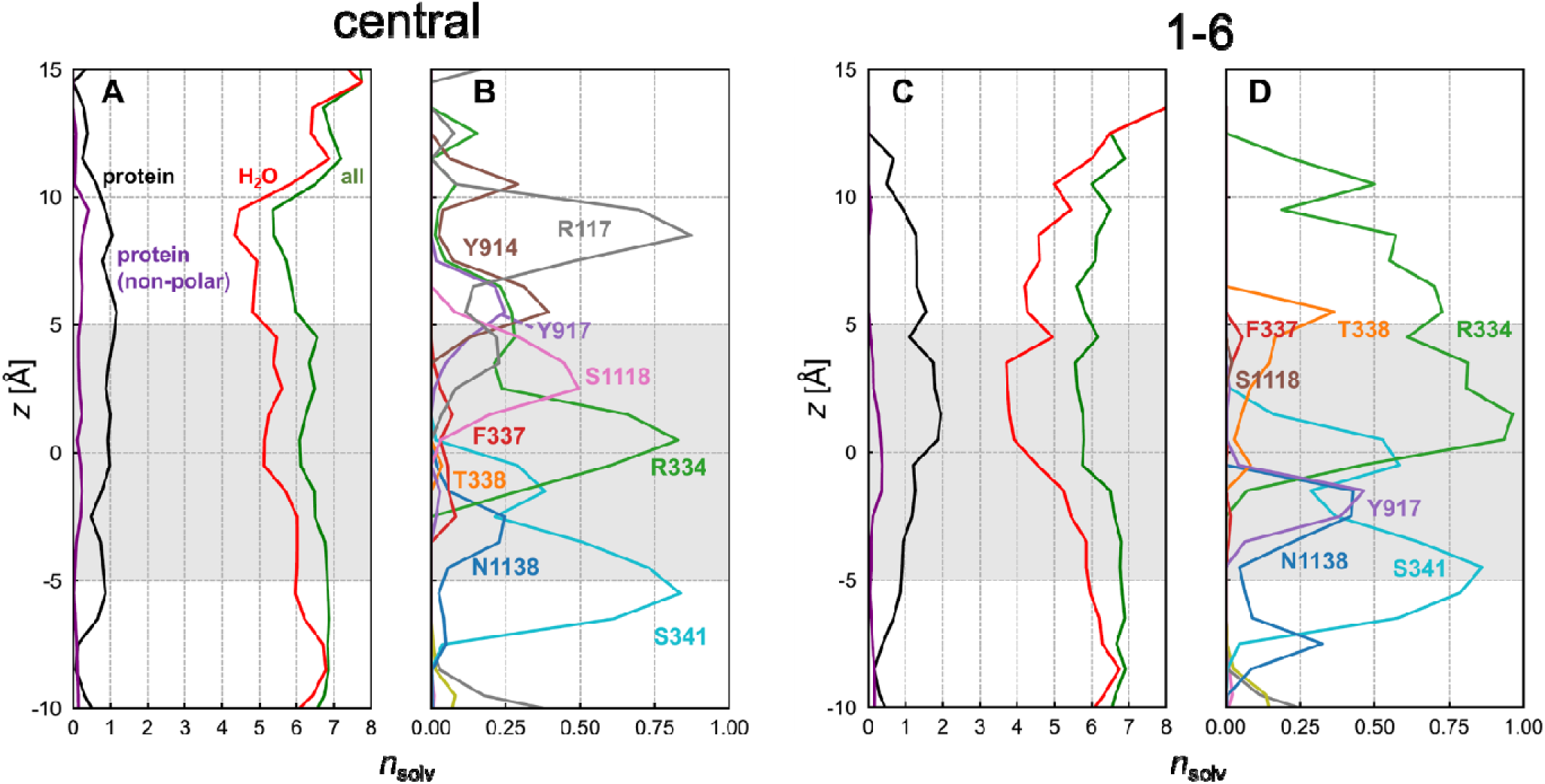
Solvation of Cl^-^ ions by permeation pathway in the pore bottleneck. Statistics of solvation by water and protein of Cl^-^ ions that followed **(A-B)** the 1-6 pathway and **(C-D)** the central pathway (combined 1-12 and intermediate pathways) in the pore bottleneck. The grey shaded region indicates the pore bottleneck (-5 < *z* < 5 Å). For details, see the caption of Figure 6.

**Figure 8 – figure supplement 1.**
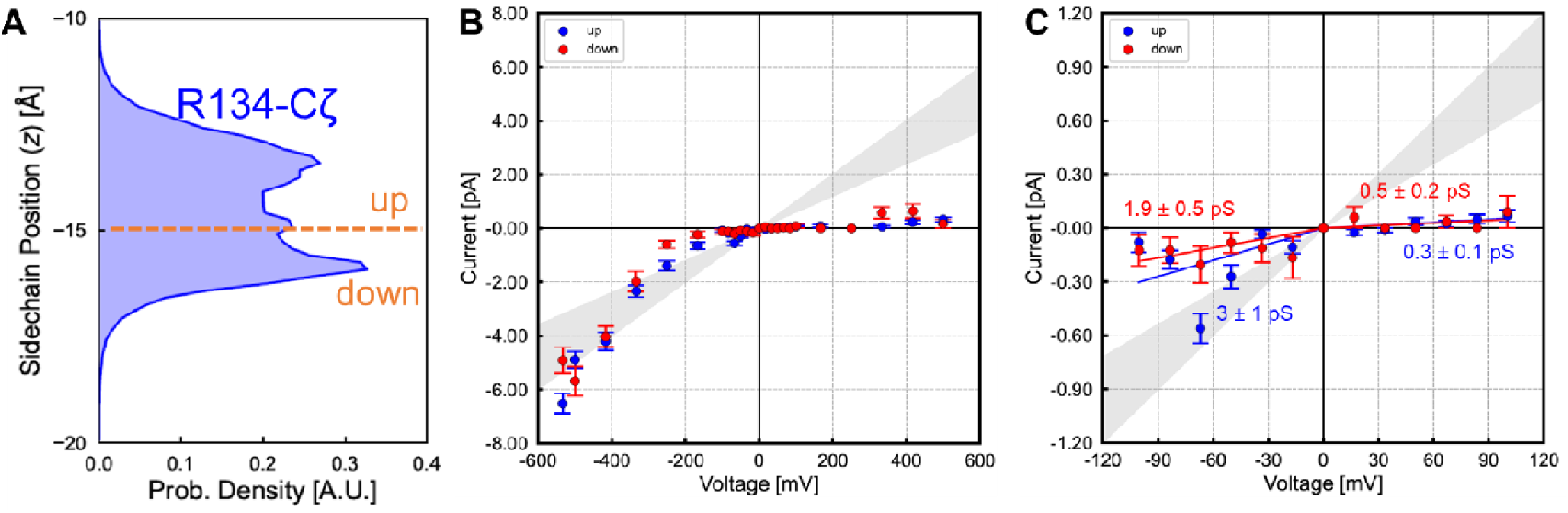
The sidechain conformation of R134 does not influence Cl^-^ permeation in CFTR. All analyses are restricted to the α state. **(A)** The axial position of the centre of charge (Cζ atom) of the R134 sidechain samples two main states “up” and “down”. **(B-C)** Estimates of Cl^-^ conductance computed from these two conformational states do not differ significantly.

**Figure 8 – figure supplement 2.**
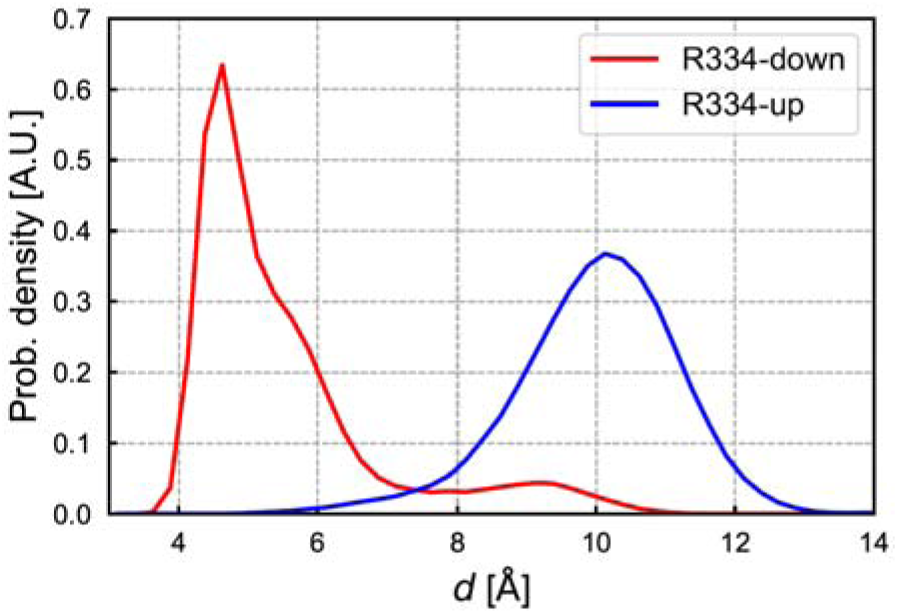
Distribution of distance (*d*) between centre of charge (Cζ atom) of R334 and centre of the phenyl ring of F337 in conformational states R334-up (blue) and R334-down (red).

**Figure 9 – figure supplement 1.**
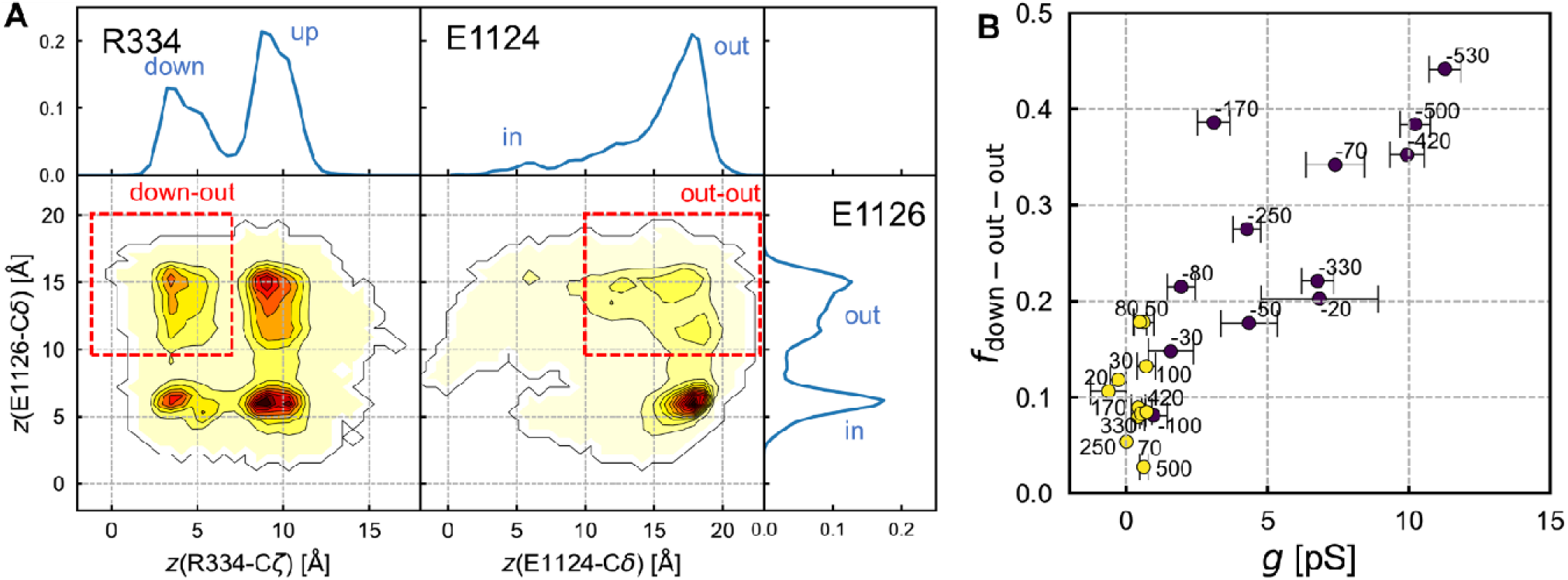
Analysis of charged sidechain arrangements in the extracellular mouth of the penta-helical pore (state α). **(A)** Axial distribution of the centres of charge of R334 (Cζ atom), E1124 and E1126 (Cδ atoms). Ion pairing between R334 and either glutamate residue is precluded when glutamate sidechains are oriented outwards. In the R334 vs E1126 contour plot (bottom left panel), the boxed region highlights the “down-out” state with R334 down and E1126 out. In the E1124 vs E1124 contour plot (bottom middle panel), the boxed region highlights the “out-out” state with both glutamate residues out. **(B)** Scatter plot of apparent Cl^-^ conductance (*g*) versus fractional population of R334 down without salt-bridge with ECL-6 glutamate residues [*f*_down-out-out_, i.e. intersection of the two red boxes in **(A)**] at different TM voltages (indicated in mV in black). The circles are colour coded according to the sign of the voltage (yellow: positive; black: negative). The positive correlation shows that at negative voltages, the sidechain conformations of the ECL-6 loop glutamate residues and R334 modulate Cl^-^ conductance. At positive voltages, the systematic under-sampling of the R334-down and glutamates-out conformation provides a plausible explanation for the apparent inward rectification of CFTR in the simulations.

**Figure 10 – figure supplement 1.**
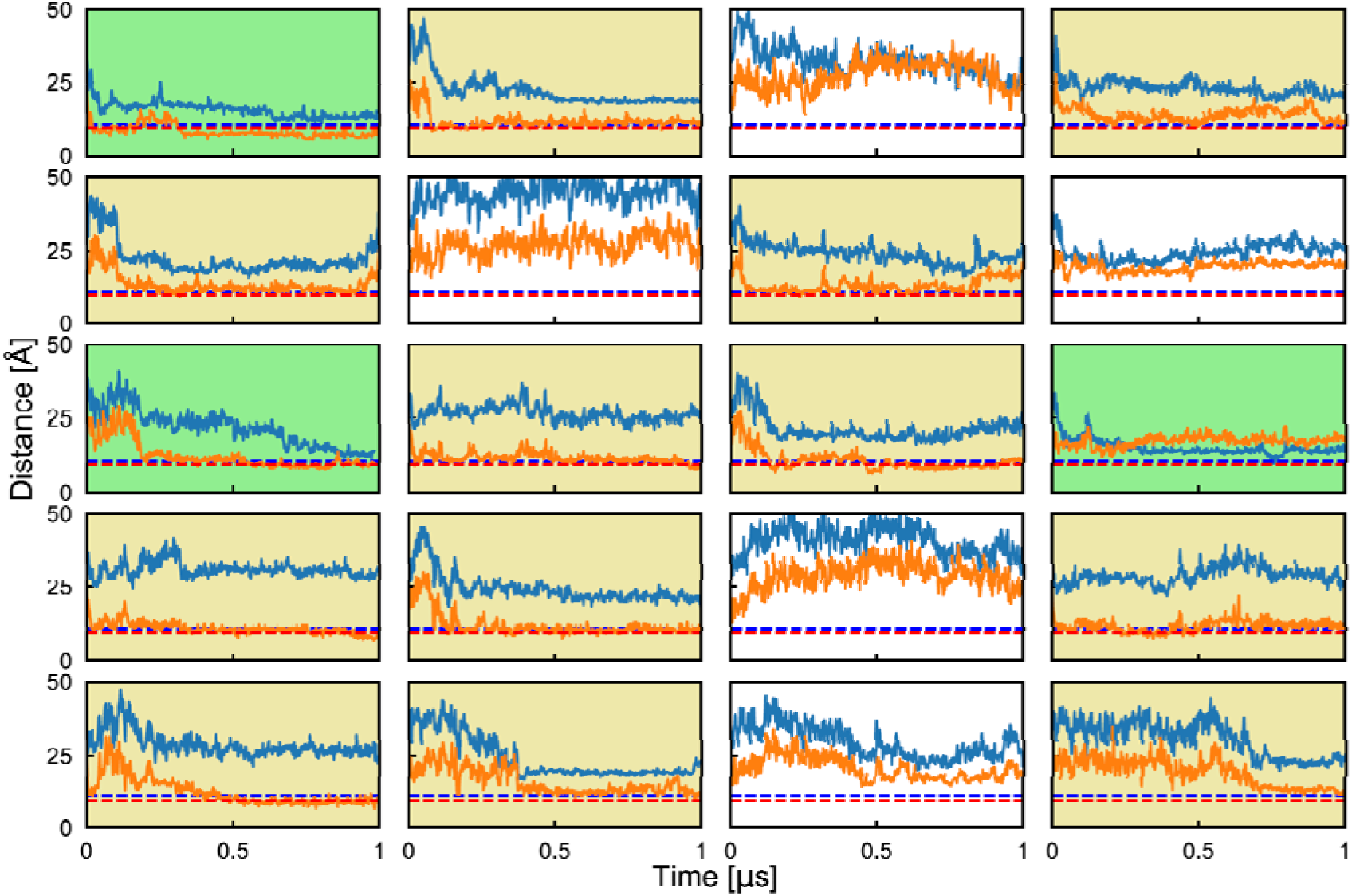
NBD1-NBD2 separation in the 20 simulations of ATP-free CFTR. Time evolution of the distances separating the two halves of the catalytic site (orange) and of the degenerate site (blue). Dashed lines indicate the distances in ATP-bound, NBD-dimerized PDB structure 6MSM (red: catalytic site; blue: degenerate site). Background colors highlight the simulations in which (green) full dimerization; (caramel) partial dimerization; or (white) no dimerization occurred.

**Figure 10 – figure supplement 2.**
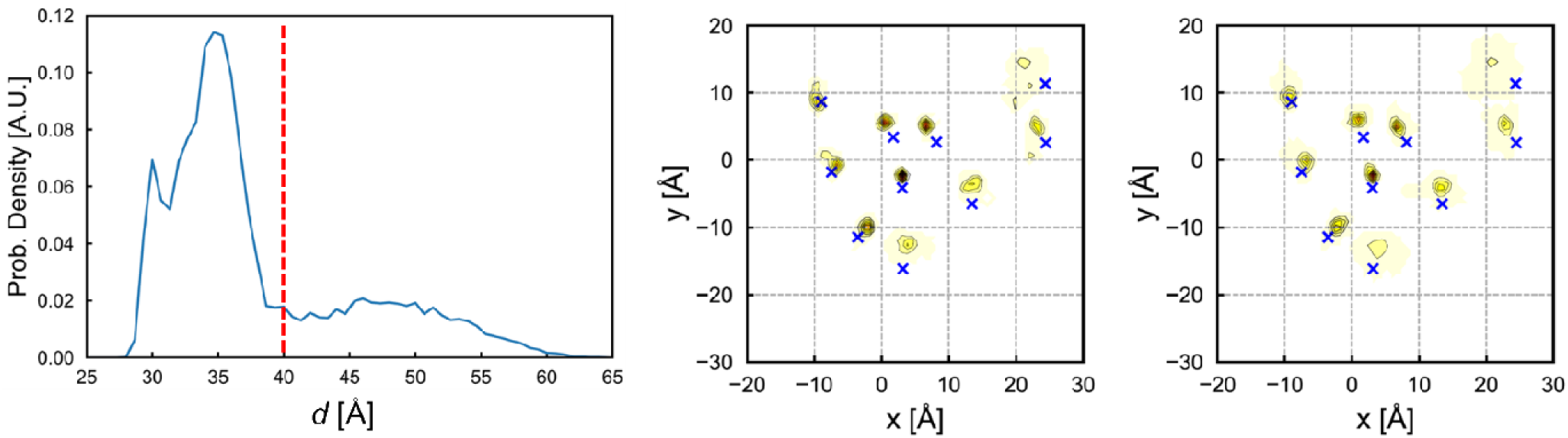
Conservation of TM helix arrangement at the extracellular end of the pore despite fast NBD dimerization in simulations of ATP-free, closed state CFTR. (Left) Distribution of centre-of-mass distance between NBDs showing an NBD-separated population (*d* > 40 Å) and NBD-dimerized population (*d* < 40 Å). Distributions of (x,y) positions of TM helices at the extracellular end **(middle)** before and **(right)** after NBD-dimerization show that their relative arrangement is conserved. Blue crosses in the 2D contour plots indicate the position of TM helices in the ATP-free, unphosphorylated, and NBD-separated human CFTR (PDB: 5UAK) at the start of the simulations.

**Figure 10 – video 1. Conformational changes from ATP-free closed state to open state illustrated by linear morph between the two states. Top view of the pore.** Opening of the channel involves reorganization of pore helices TM 12 (yellow) and 8 (green) relative to TM 1 (blue), 2 (red), 6 (tan), and 11 (magenta). TM11 separates from TM10 (cyan) to accommodate TM12.

**Figure 11 – figure supplement 1.**
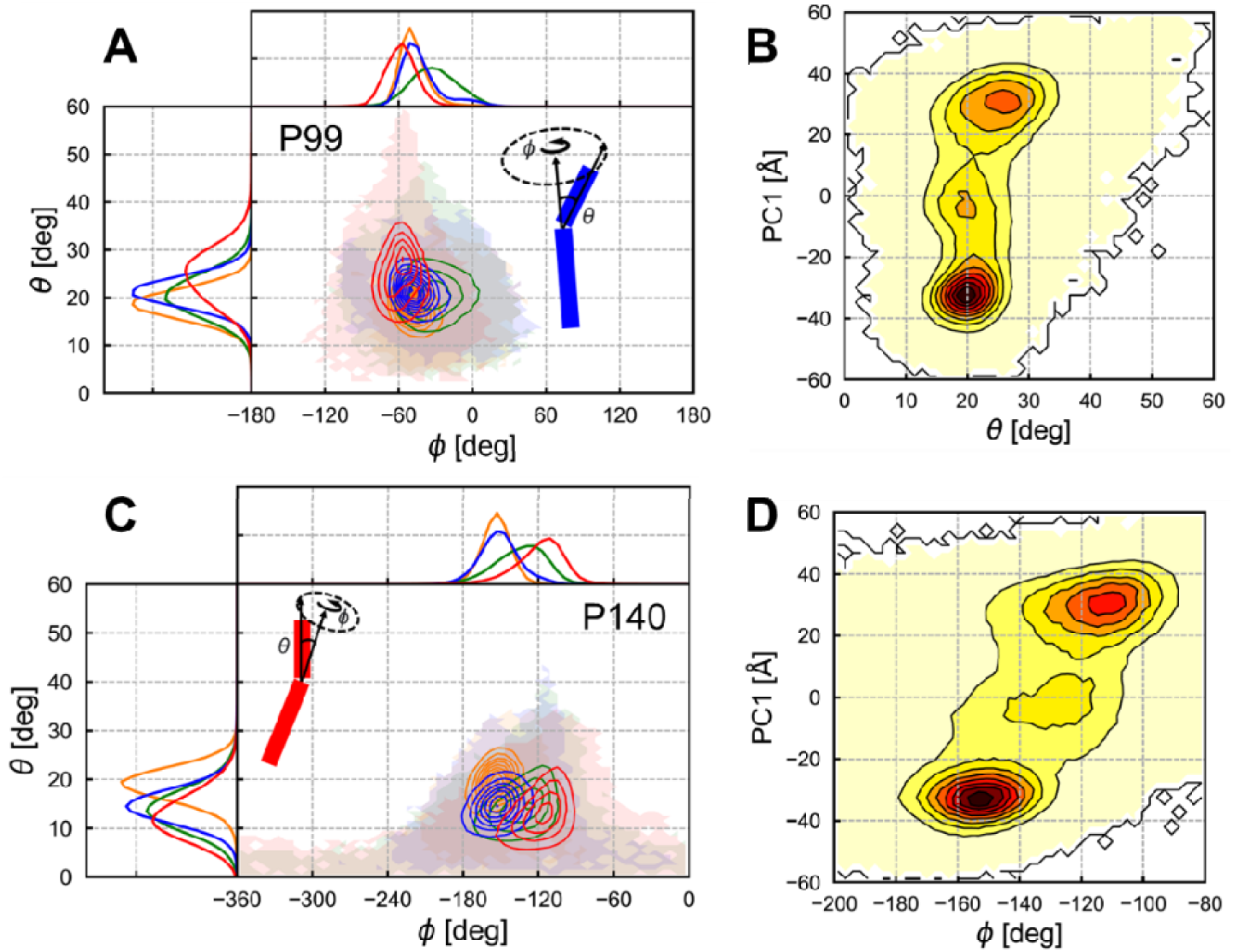
Analysis of helical kink and orientation angles at **(A-B)** P99 on TM1 and **(C-D)** P140 on TM2. See the caption of Fig. 11 for details.

**Figure 11 – video 1. Conformational changes from MD-closed state β to MD-open state** α visualized using 50 randomly selected snapshots from each state (first 50 frames: state β, last 50 frames: state α). Note the changes in the conformation and orientation of TM helices 1 (blue), 2 (red), and 11 (magenta) relative to TM12 (yellow) and the other pore lining helices, TM 6 and 8 (grey).

